# Gonadal PIP-seq reveals genes involved in germ cell development and sexual differentiation in sea lamprey (*Petromyzon marinus*)

**DOI:** 10.64898/2026.06.10.731425

**Authors:** Yu-Wen Chung-Davidson, Stephanie L. Hickey, Matthew P. Bernard, Sreeja Sarasamma, Ciro Amato, Martin Estermann, Jacob Kimmel, Lily Gorman, Tyler Buchinger, Humprey Yao, Weiming Li

**Affiliations:** Department of Fisheries & Wildlife, Michigan State University, East Lansing, MI 48824; Bioinformatics Core Facility, Michigan State University, East Lansing, MI 48824; Department of Pharmacology & Toxicology, Michigan State University, East Lansing, MI 48824; Program of Environmental Biology & Plant Biology, Michigan State University, East Lansing, MI 48824; Reproductive Developmental Biology Group, National Institute of Environmental Health Sciences, Research Triangle Park, NC 287709

## Abstract

Vertebrate sex determination is highly diverse, with distinct mechanisms that have evolved repeatedly and independently across phylogeny. The sea lamprey (*Petromyzon marinus*) is an extant jawless vertebrate that remains sexually labile during its larval stage that lasts around 3-20 years. Their gonad contains bipotential, female, and male germ cells before metamorphosis. To examine the genetic programs in germ cells at different developmental stages, we profiled transcriptomes of dissociated gonadal cells from larvae and recently metamorphosed juvenile (transformers, sex-determined) using fluorescence-activated cell sorting (BD FACS Aria IIu) and PIPseq^TM^ (Fluent BioSciences/Illumina). 66,676 cells were sequenced and classified into 19 cell clusters according to their gene expression profiles. Primordial germ cells differentially expressed genes encoding various heat-shock proteins, hormones, growth factors, and their receptors, and proteins sensitive to iron, nitrogen starvation, sugar and lipid metabolisms, and oxidative stress, providing possible links between environmental factors, cellular functions, and cell fate determinants. Germ cells in larvae had enriched expression of genes involved in DNA replication, transcription, translation, apoptosis, autophagy, cell cycle, cell differentiation, cell adhesion and migration, cell survival, chromatin remodeling, and proliferation. Specifically, female germ cells differentially expressed various transcript isoforms of AEP1, CUZD1, S100A1, and ZP genes, whereas male germ cells differentially expressed BOLL and PIWIL1. Pseudotime trajectory and Gene Ontology (GO) analysis revealed that genes with changing expressions along the female differentiation pathway were involved in binding of sperm to zona pellucida, egg coat formation, prevention of polyspermy and positive regulation of acrosome reaction. On the other hand, genes with changing expressions along the male differentiation pathway were involved in ribosome assembly and translation. Several WNT5 transcripts were differentially expressed in germ cells or somatic cells. Our results suggest that sea lamprey sex determination and sexual differentiation likely involve the coordinated action of numerous genes that interact with various environmental factors and regulate germ cell specification and development.

## Introduction

The mechanisms of sex determination and sexual differentiation are strikingly variable among species. In contrast to highly conserved developmental processes that are regulated by the same upstream gene networks, not one single gene initiates sex determination in all species (Capel, 2017). Among vertebrates, different cell types in the gonad can initiate the process of sex determination. In mammals, somatic cells seem to induce sexual differentiation processes, whereas in fishes germ cells appear to assume the driver’s seat (Capel, 2017). During sexual differentiation, the reproductive system contains bipotential primordial cells. As a result, each embryo exhibits the potential to differentiate into either male or female (Capel, 2017). In gonochoristic species, sex determination gene(s) activates sexual differentiation pathway of one sex and inhibits the differentiation pathway for the opposite sex. In sea lamprey, female and male pathways may coexist during early gonadal development, but male pathway seems to require more genes located in the germline specific region to initiate or maintain the maleness, or both (Yasmin et al., 2022).

Sex determination systems are typically classified as genetic sex determination (GSD) or environmental sex determination (ESD). However, many vertebrate species can exert both GSD and ESD mechanisms simultaneously (Holleley, 2015; Ezaz, 2006). Sea lamprey resides at the earliest branching position of vertebrate phylogenetic tree (Delsuc et al., 2006), offering a valuable opportunity to investigate the diversity and evolution of vertebrate sex determination mechanisms.

Lampreys are characterized by a long larval period with undifferentiated gonads (Okkelberg 1921; Hardisty 1965). After metamorphosis, lampreys commit to a specific sex (Lowartz & Beamish, 2000). Parasitic lampreys, those species that have retained the juvenile period during which they feed on the body fluids and tissues of fishes, usually develop a large number of oocytes in the gonad (Beamish & Potter, 1975). Although the mechanisms underlying sex determination in lamprey species are unclear, environmental factors such as population density and growth rate are suspected to bias the sex ratio in the population (Johnson et al., 2017). The proportion of males increased significantly with relative population density (Docker & Beamish, 1994). Under conditions promoting rapid growth, the percentage of males varied directly with density and inversely with temperature. However, temperature had little effect when larval growth was slow (Beamish, 1993).

To uncover the gene expression pathway associated with primordial, male and female germs cells in sea lamprey, we used Particle-templated Instant Partition sequencing (PIP-seq), a microfluidics-free single-cell RNA-sequencing technique to examine dissociated gonadal cells from sex-undetermined larvae and sex-committed female and male transformers. Subsequent pseudotime trajectory analysis of germ cell gene expression profiles revealed the genetic program and pathways specific for each sex in sea lamprey.

## Methods and Materials

### Animals

Larval (sex-labile; body length: 11.2 ± 0.1 cm, body weight: 2.0 ± 0.0 g) and newly transformed (sex-determined, referred as transformers; body length 13.1 ± 0.1 cm, body weight: 1.9 ± 0.1 g) sea lamprey for PIP-seq and extra animals in different developmental stages (larvae, body length: 9.2 ± 0.1 cm, body weight 1.2 ± 0.0 g; transformers, body length: 12.6 ± 0.3 cm, body weight: 5.9 ± 0.1 g; adult sea lamprey, body length: 50.3 ± 2.0 cm, body weight: 211.0 ± 38.0 g) were provided by the staff of the U.S. Geological Survey, Hammond Bay Biological Station, Great Lakes Science Center, Millersburg, MI, U.S.A. All animals were transferred to Michigan State University where standard operating procedures for transporting, maintaining, handling, anesthetizing, and euthanizing sea lampreys were approved by the Institutional Committee on Animal Use and Care of Michigan State University and in compliance with standards defined by the National Institutes of Health Guide for the Care and Use of Laboratory Animals (Institute for Laboratory Animal Research, 2011). All applicable international, national, and/or institutional guidelines for the care and use of animals were followed.

### Gonadal cell dissociation

Six larvae and three transformers of each sex were euthanized with 0.65 mg/ml ethyl 3-aminobenzoate methanesulfonate (MS222, Sigma-Aldrich Co. LLC, St. Louis, MO, U.S.A.). Gonadal tissues were dissected out and placed in 5 ml propylene tube containing 3 ml HBSS (ThermoFisher; Life Technologies Corporation, Carlsbad, CA, U.S.A.) on ice (3 gonads/tube). Gonadal cells were dissociated using a ThermoScientific^TM^ Pierce^TM^ Primary Neuron Isolation Kit (ThermoFisher No. 88280) according to the manufacturer’s instructions. Briefly, gonadal tissues were rinsed with 4 ml HBSS, then incubated in 1ml neuronal isolation enzyme with papain reconstituted in HBSS in an incubator shaker (15 rpm) at room temperature for 30 min. Dissociated gonadal cells were then centrifuge at 200 x g, 4°C for 3 min. The supernatant was gently decanted, and 1 ml HBSS was added to the pellet and centrifuged at 200 x g, 4°C for 3 min. The supernatant was gently decanted, and 500 µl of serum supplemented Neuronal Culture Media (10% fetal bovine serum, 1x glutamine supplement, and 1% penicillin/streptomycin) was added to the tissue pellet. The enzyme-digested gonadal tissues were gently broken down by gently pipetting up and down 20 times using a 1000 µl wide-bore pipette tip set at 400 µl. After that, 500 µl of serum supplemented Neuronal Culture Media was added to the suspension, and the suspension was pressed through a 40 µm BelArt^TM^ Flowmi^TM^ tip strainer (ThermoFisher). Cell concentration and cell viability were examined use a BioRad TC20 Automated Cell Counter using 10 µl of cell suspension with 10 µl trypan blue mixture (BioRad Laboratories, Corp., Hercules, CA, U.S.A.). Cell suspension was centrifuged at 200 x g, 4°C for 3 min, the supernatant was discarded, and flow cytometry cell sorting buffer (HBSS with 1% heat-inactivated fetal bovine serum, 1 mM EDTA, and 25 mM HEPES, pH 7.0) was added to the pellet to adjust the cell concentration to around 1 x 10^6^ cells/ml and split into two 5 ml propylene tubes. One tube served as the unstained control, and the other tube was added with 2 µl/ml 7-aminoactinomycin D (7-AAD, ThermoFisher) for Fluorescence-activated Cell Sorting (FACS) using a BD FACS Aria IIu (BD Biosciences, San Jose, CA, U.S.A.). Live cells (7-AAD negative) were collected in a 5 ml propylene tube coated with 500 µl fetal bovine serum.

### PIP-seq

Samples were prepared using the PIPseq™ T20 3ʹ Single Cell Capture and Lysis Kit v4.0 following the instructions provided by Fluent Biosciences (now part of Illumina, Inc., San Diego, CA, U.S.A.). Briefly, FACS collected cells were centrifuged at 200 x g, 4°C for 3 min. The supernatant was discarded, and 1 ml Cell Suspension Buffer was added and gently mixed 5 times using a P1000 wide-bore low retention pipette tip. The cell suspension was centrifuged at 200 x g, 4°C for 3 min, and the supernatant was discarded. 400 µl Cell Suspension Buffer was added to the pellet and gently mixed 15 times using a standard-bore P1000 pipette tip to create a homogeneous cell suspension. The cell suspension was pressed through a 40 µm Flowmi tip cell strainer. Cell concentration and viability was determined using a BioRad TC20 automated cell counter. Cell concentration was adjusted to 4 x 10^6^ cells/ml, and 10 µl of the cell suspension was added to the PIPs and mixed 10 times with a P200 pipette set to 180 µl. 1000 µl partitioning reagent was added to the cell/PIP mixture and vortex horizontally at 2000 rpm for 15 sec, and vortex vertically at 2000 rpm for 2 min using the Starter kit equipment (Fluent BioSciences/Illumina). The PIP tubes were sat on a provided stand for 30 seconds, and extra partitioning reagents were removed using a provided syringe with G22 blunt tip needle. 630 µl chemical lysis buffer emulsion was added to the top of the PIP emulsion, and the emulsion was incubated in the dry bath provided in the Starter kit using Program A and shipped to the company for further processing (library preparation, sequencing, and data analysis). Four samples were submitted: C00004336 (larval duplicate 1, submitted on July 11, 2023), C00004337 (female transformer, submitted on November 8, 2023), C00003262 (larval duplicate 2, submitted on July 11, 2023), and C00004374 (male transformer, submitted on November 8, 2023). Data were deposited in National Center for Biotechnology Information (NCBI) Gene Expression Omnibus (GEO) (https://www.ncbi.nlm.nih.gov/geo/) under GSE334273.

### Data analysis

All analyses were performed using R Statistical Software (v4.4.1) (R Core Team, 2021).

### Quality control

Quality control steps were performed for each sample separately. To identify genuine cells from empty droplets, the emptyDrops function from the *DropletUtils* (v1.26.0) package was used, setting the by.rank parameter to 20K cells (Lun et al., 2019). Barcodes with an FDR < 0.001 were retained as likely cell-containing droplets. Quality control metrics were computed using perCellQCMetrics from the *scater* (v1.34.1) package (McCarthy et al., 2017). Mitochondrial genes were identified as those on the mitochondrial chromosome (NC_001626.1). Cells with percent of mitochondrial transcripts > 3 median absolute deviations (MAD) above the median percent mitochondrial transcripts, as calculated by the isOutlier function from the package, were removed (McCarthy et al., 2017).

### Normalization and clustering

Normalization steps were performed using the *scran* (v1.34.0) package (Lun et al., 2016). Counts from each sample underwent log-normalization using logNormCounts. Feature selection was performed on each dataset individually by modeling gene variances with modelGeneVar, followed by selecting the top 10% highly variable genes (HVGs) via getTopHVGs. To merge individual datasets and correct for batch effects, a common set of genes were identified across all samples via intersection. Using this gene universe, expression matrices and variance statistics for each sample were subset to include only shared genes. To account for depth differences, we applied multiBatchNorm from the *batchelor* (v1.22.0) package (Haghverdi et al., 2018) across the samples. HVGs were then re-identified using the combined variance model generated via combineVar from the *scran* package (v1.34.0), and the top 5,000 HVGs were retained (Lun et al., 2016). Sample integration was conducted using the fastMNN method from the *batchelor* (v1.22.0) package, using the top HVGs as input features (Haghverdi et al., 2018). The resulting merged dataset, containing batch-corrected PCA embeddings, was clustered using a shared nearest neighbor graph (k = 30) followed by Louvain clustering.

### Identification of cluster enriched genes

To identify enriched genes for each cell cluster in the integrated dataset, we performed differential expression (DE) analysis using the findMarkers function from the *scran* (v1.34.0) package using each sample’s normalized log-expression values (Lun et al., 2016). A two-sample t-test was used for each gene to pairwise compare expression in each cluster against all others. To account for inter-sample variability, we applied blocking by sample origin. The direction="up" parameter was used to focus on genes that were upregulated in each cluster compared to others. The pval.type="some" option was used to retain genes with statistically significant differential expression in some but not necessarily all pairwise comparisons, enhancing sensitivity in heterogeneous datasets. Clusters were annotated by comparing cluster enriched genes with cell-type marker genes collected from the literature (Supplementary Table S1).

### Gene Ontology Annotation

To assist in annotating cell type clusters, we performed Gene Ontology (GO) enrichment analysis on the top 100 marker genes for each cluster using a hypergeometric test for overlap with the total number of assayed genes as the gene universe (21,967). GO annotations for *Petromyzon marinus* genes were retrieved from https://ftp.ncbi.nlm.nih.gov/genomes/all/GCF/010/993/605/GCF_010993605.1_kPetMar1.pri/GCF_010993605.1_kPetMar1.pri_gene_ontology.gaf.gz. The annotations were preprocessed by filtering out entries with the “NOT” qualifier to remove explicitly negated annotations. To reduce noise and avoid overly broad or overly narrow terms, we filtered GO terms to include only those associated with more than 10 and fewer than 200 genes. False discovery rates (FDR) were calculated using the Benjamini-Hochberg correction. Only GO terms with p-value < 0.01 were retained for downstream analysis and cluster annotation.

### Germ cell subclustering

Cells from the following germ cell clusters were selected from the full data set: cluster 5 (oogonia/oocytes), cluster 7 (migrating primordial germ cells), cluster 16 (cytokinetic spermatocytes), cluster 17 (migrating male germ cells), cluster 18 (apoptotic spermatocytes), and cluster 19 (GPCR-activated spermatocytes). The raw counts were renormalized and subclustered using the same methods as the full dataset. Briefly, normalization and variance modeling were carried out with *scran* v1.34.0 (Lun et al., 2016), and batch correction and integration were performed using *batchelor* v1.22.0 via fastMNN (Haghverdi et al., 2018). Clustering was based on a shared nearest neighbor graph (k = 8) and Louvain algorithm. Cluster-enriched genes were identified with findMarkers (*scran*), blocking by sample (Lun et al., 2016).

### Trajectory analysis

We conducted a pseudotime analysis on integrated germ-cell RNA-seq data using the *TSCAN* (v1.44.0) package framework (Ji & Ji, 2017). Pseudotime was inferred using quickPseudotime applied to the integrated dataset. Specifically, a minimum spanning tree was constructed based on cluster centroids computed in the corrected dimensionality-reduced space from MNN integration. Pseudotemporal ordering was assigned to cells starting from a specified root node (cluster 2 primordial germ cells in this case). We computed a consensus pseudotime ordering for visualization using averagePseudotime. Statistical testing of gene expression along each branch of the pseudotime trajectory was performed with the testPseudotime function using each sample’s normalized log-expression values. Gene ontology enrichment analyses were performed as described above on the top 100 up- and down-regulated genes by FDR along each path, as well as the male and female branch-point-specific genes.

### LOC gene functional annotation

Human homologs or homologs from other species derived by Wang et al. (2024) were used to further annotate genes with LOC numbers in the NCBI sea lamprey reference genome (kPetMar1.pri) when possible. See Supplementary Table S2 for a map from kPetMar1 gene IDs to Wang et al. gene IDs.

### Immunohistochemistry and Immunofluorescent Staining

Sea lamprey gonadal tissues were fixed in 4% paraformaldehyde in 0.1M phosphate buffer (pH 7.4) for at least 24 h, rinsed with 0.1M phosphate buffer saline (PBS, pH 7.4) for 5 min, dehydrated in increasing ethanol series (10%∼70% ethanol with 10% increment each for 5 min), and sent to Michigan State University Investigative Histopathology Laboratory to process for paraffin sections. Paraffin sections were deparaffinized in xylene for 5 min 3 times, rehydrated through 100% ethanol for 5 min, 95% ethanol for 5 min, 70% ethanol for 5 min, and PBS for 5 min twice.

Immunohistochemistry staining was conducted according to the manufacturer’s instruction. Briefly, gonadal sections (on the slides) were incubated with BLOXALL Endogenous Blocking Solution (Vector Laboratories, Inc., Newark, CA, U.S.A.) for 10 min and washed in PBS for 5 min twice to eliminate endogenous peroxidases and phosphatases. Gonadal sections were then incubated in Avidin solution (Avidin-Biotin Blocking Kit, Vector) for 15 min, washed with PBS for 5 min, and incubated in Biotin solution (Avidin-Biotin Blocking Kit, Vector) for 15 min to eliminate endogenous biotin. After PBS wash for 5 min, the sections were incubated in primary antibody with normal goat serum (VECTASTAIN Elite ABC-HRP Kit, Vector) in PBS with 0.1% Triton X-100 (Sigma) at 4°C overnight. The sections were then washed with PBS for 5 min 3 times, incubated in biotinylated goat-anti-rabbit secondary antibody with normal goat serum (VECTASTAIN Elite ABC-HRP Kit, Vector) in PBS with 0.1% Triton X-100 for 30 min at room temperature, and washed in PBS for 5 min 3 times. The sections were incubated in ABC solution (VECTASTAIN Elite ABC-HRP Kit, Vector) for 30 min and washed with PBS for 5 min 3 times, and subsequently incubated in ImmPACT DAB (ImmPACT DAB Peroxidase Substrate kit, Vector) for 2-10 min, washed in tap water for 5 min 2 times, and counter stained with Hematoxylin QS (Vector) for 45 sec. Gondal sections were washed with tap water, dehydrate through 70%, 95%, and 100% ethanol, 5 min each. The stained sections were clarified in xylene for 5 min 3 times, coverslipped with DPX mounting media (Sigma), and imaged using an ECHO Revolution imaging system (ECHO, a BICO Company, San Diego, CA, U.S.A.). Primary antibodies: 1:100 rabbit-anti-BOLL (HPA018678, Sigma); 1:500 rabbit-anti-CXCR4 (SAB3500383, Sigma); 1:100 rabbit-anti-FOXL2 (F0005, Sigma); 1:1000 rabbit-anti-RBPMS2 (SAB2106506, Sigma); 1:1000 rabbit-anti-S100A1 (HPA006462, Sigma); 1:250 rabbit-anti-SOX18 (SAB2108480, Sigma).

For immunofluorescent staining, paraffin sections were deparaffinized, rehydrated, and incubated in primary antibody as described above, and then incubated in fluorescent secondary antibody, and coverslipped using ProLong Diamond antifade mounting media with DAPI (ThermoFisher). Primary antibodies: 10 µg/ml rabbit-anti-DDX4 (Invitrogen PA5-23378, ThermoFisher); 0.85 µg/ml rabbit-anti-PIWIL1 (Invitrogen PA5-17034, ThermoFisher); 10 µg/ml rabbit-anti-SYCP3 (Bio-techne/Asuragen, Inc., Austin, TX, U.S.A.); 0.5 µg/ml rabbit-anti-WNT5B (Invitrogen PA5-33114, ThermoFisher). Secondary antibodies: 1 µg/ml goat-anti-rabbit IgG-Alexa 594 (ThermoFisher).

### HCR™GoldRNA-FISH (HCR)

Marker gene mRNA was detected using designed probes from Molecular Instruments, Inc. (Los Angeles, CA, U.S.A.) using mRNA sequence retrieved from the NCBI sea lamprey genome website (https://www.ncbi.nlm.nih.gov/datasets/genome/GCF_010993605.1/). The hybridization procedure followed the manufacturer’s instruction. Briefly, paraffin sections were prepared as described in Immunohistochemistry and Immunofluorescent Staining section. Wash buffer used PBS containing 0.1% diethylpyrocarbonate (DEPC) and 0.3% Tween 20 (PBST-DEPC). Gonadal sections were treated with 5 µg/ml proteinase K in PBST-DEPC at room temperature for 10 min, washed with PBST-DEPC for 5 min 3 times. Gonadal sections were prehybridized in a humidifying chamber with HCR^TM^ HiFi Probe Hybridization Buffer at 37°C for 10 min and hybridized with the designed probe mix (detail see below) in HCR^TM^ HiFi Probe Hybridization Buffer in a humidifying chamber at 37°C overnight. Gondal sections were washed with HCR™ HiFi Probe Wash Buffer at 37°C for 15 min 4 times, incubated in HCR™GoldAmplifier Buffer at room temperature for 30 min in a humidified chamber, and continued incubated with fluorescent hair pin amplifier solution at room temperature overnight in the dark. Gonadal sections were washed with HCR™GoldAmplifier Wash Buffer at room temperature for 15 min 4 times, 5x SSC at room temperature for 15 min 4 times, PBS for 15 min, coverslipped with ProLong Diamond antifade mounting media with DAPI (ThermoFisher) and imaged using a Leica Stellaris 8 DIVE imaging system (Leica Microsystems, Inc., Deerfield, IL, U.S.A.). Probe Mix 1: CD9-X1_Alexa 488; GLG1-X2_Alexa 546; HSPA5-X3_Alexa 647; PIWIL1-X8_Alexa 594. Probe Mix 2: BOLL-X1_Alexa 488; GLG1-X2_Alexa 546; TEX11-X3_Alexa 647; PIWIL1-X8_Alexa 594.

## Results and Discussion

### PIP-seq identifies germ and somatic cell types in sex-labile and sex-determined sea lamprey gonads

After data processing and quality filtering, PIP-seq captured 17,186 cells from C00004336 (larval duplicate 1), 17,454 cells from C00004362 (larval duplicate 2), 13,803 cells from C00004337 (female transformer), and 18,324 cells from C00004374 (male transformer) (Supplementary Table S3). The four datasets were merged and corrected for sample batch effects, yielding 19 cell clusters (Figure 1). We assigned cluster identities using the expression of established cell-type markers (Supplementary Table S1), genes enriched within each cluster (Supplementary Table S4), and Gene Ontology (GO) terms overrepresented among those enriched genes (Supplementary Table S5). Several clusters differentially expressed distinctive gene markers (Table 1); for example, multiple CUZD1 isoforms were expressed in the oogonia/oocyte cluster (cluster 5), but one transcript (CUZD1.15: LOC116956112) was especially enriched in this cluster (Supplementary Figure S6).

**Figure 1.**
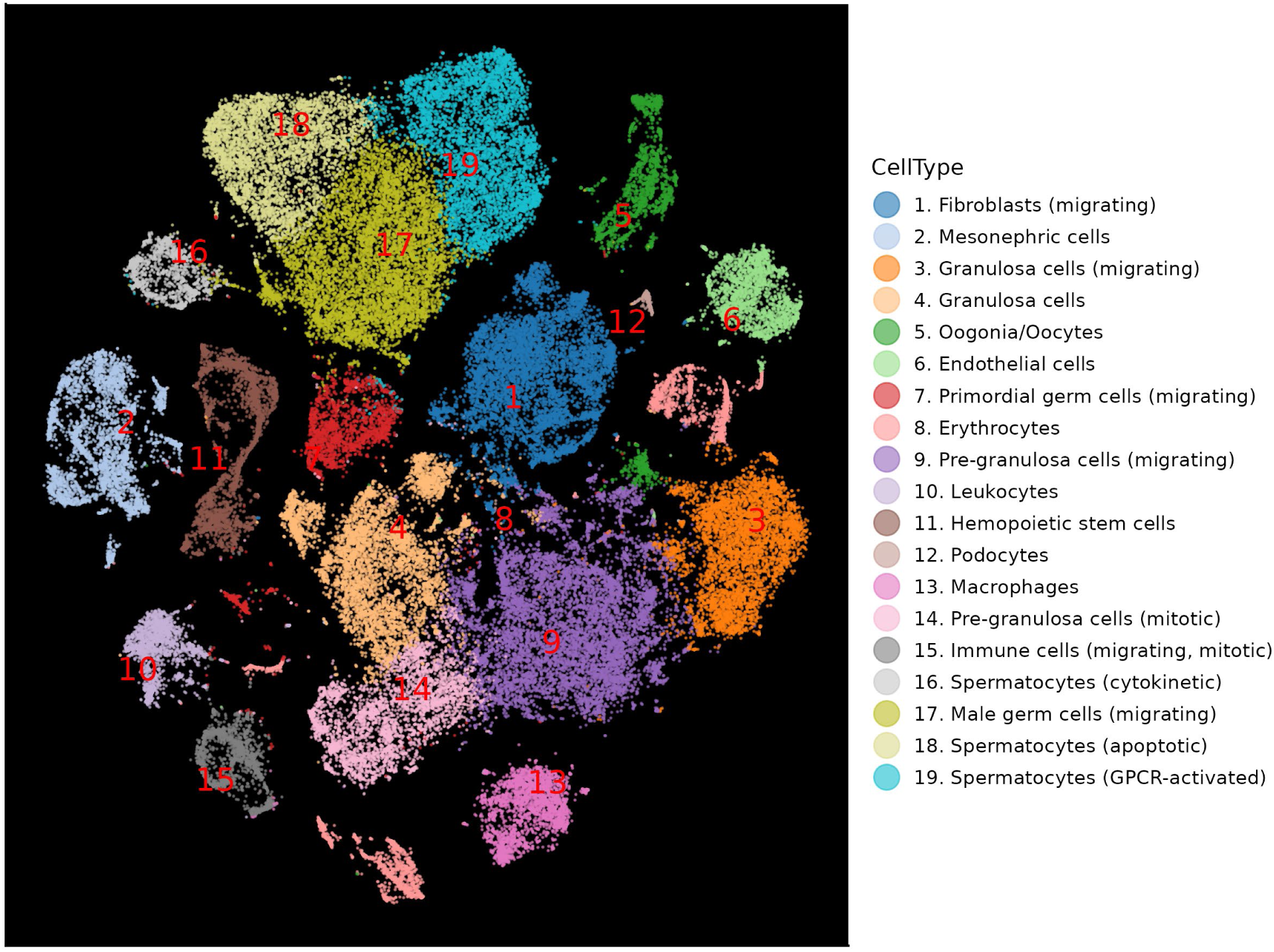
Cell clusters identified using the “scatter” package in the batch corrected TSNE expression space.

To validate these assignments, we examined selected markers at both the protein level (immunohistochemistry and immunofluorescent staining) and the transcript level (HCR™ GoldRNA-FISH) in gonadal tissues, and in each case the staining patterns were consistent with the cluster identities. DDX4, a germ cell marker enriched in clusters 5 (oogonia/oocytes) and 14 (pre-granulosa cells), produced immunoreactivity in migrating primordial germ cells, oogonia, oocytes, pre-granulosa cells, and follicular cells (Figure 2 & Supplementary Figure S7). PIWIL1, another germ cell marker, was enriched in clusters 5 (oogonia/oocytes), 7 (migrating primordial germ cells), 9 (migrating pre-granulosa cells), 16 (cytokinetic spermatocytes), 17 (migrating male germ cells), 18 (apoptotic spermatocytes), and 19 (GPCR-activated spermatocytes). PIWIL1-immunoreactivity was observed in migrating cells (primordial, female, or male germ cells) and within the germ cell cyst (oogonia or spermatogonia) (Figure 3 & Supplementary Figure S8), with HCR-FISH yielding concordant results (Figures 4 and 5). Likewise, SYCP3 was enriched in clusters 5, 7, 16, 17, 18, and 19, and SYCP3-immunoreactivity was found in migrating primordial germ cells, within the germ cell cyst, and in spermatocytes (Figure 6 & Supplementary Figure S9). Immunohistochemistry for BOLL, CXCR4, FOXL2, RBPMS, S100A1, and SOX18 also confirmed the PIP-seq results (Supplementary Figures S10 & S11).

**Figure 2.**
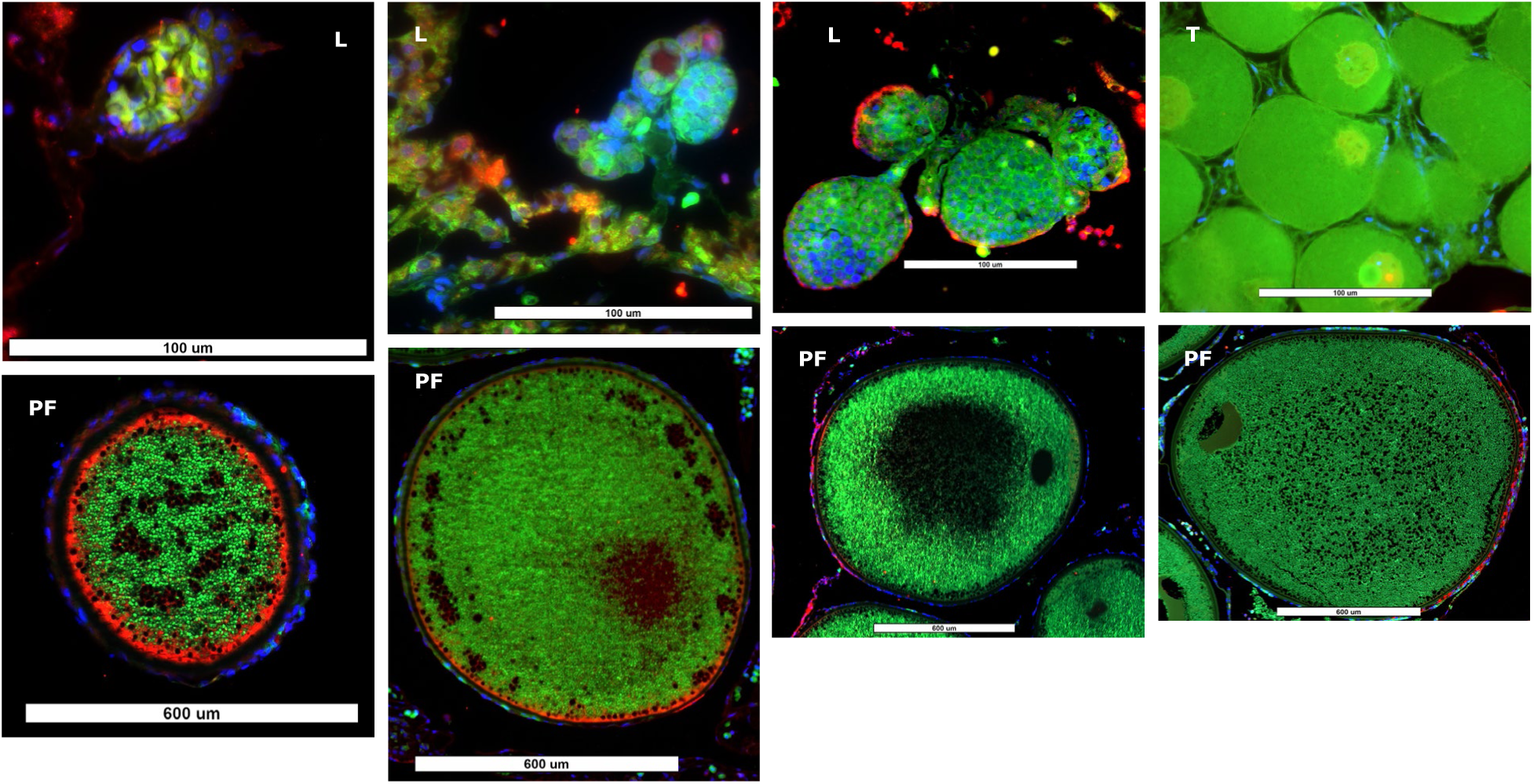
DDX4/VASA-immunoreactivities (red) in primordial germ cells, oogonia, oocytes, pre-granulosa cells, and follicular cells. Blue: DAPI; Green: autofluorescence; L: larva; PF: preovulatory female; T: transformer.

**Figure 3.**
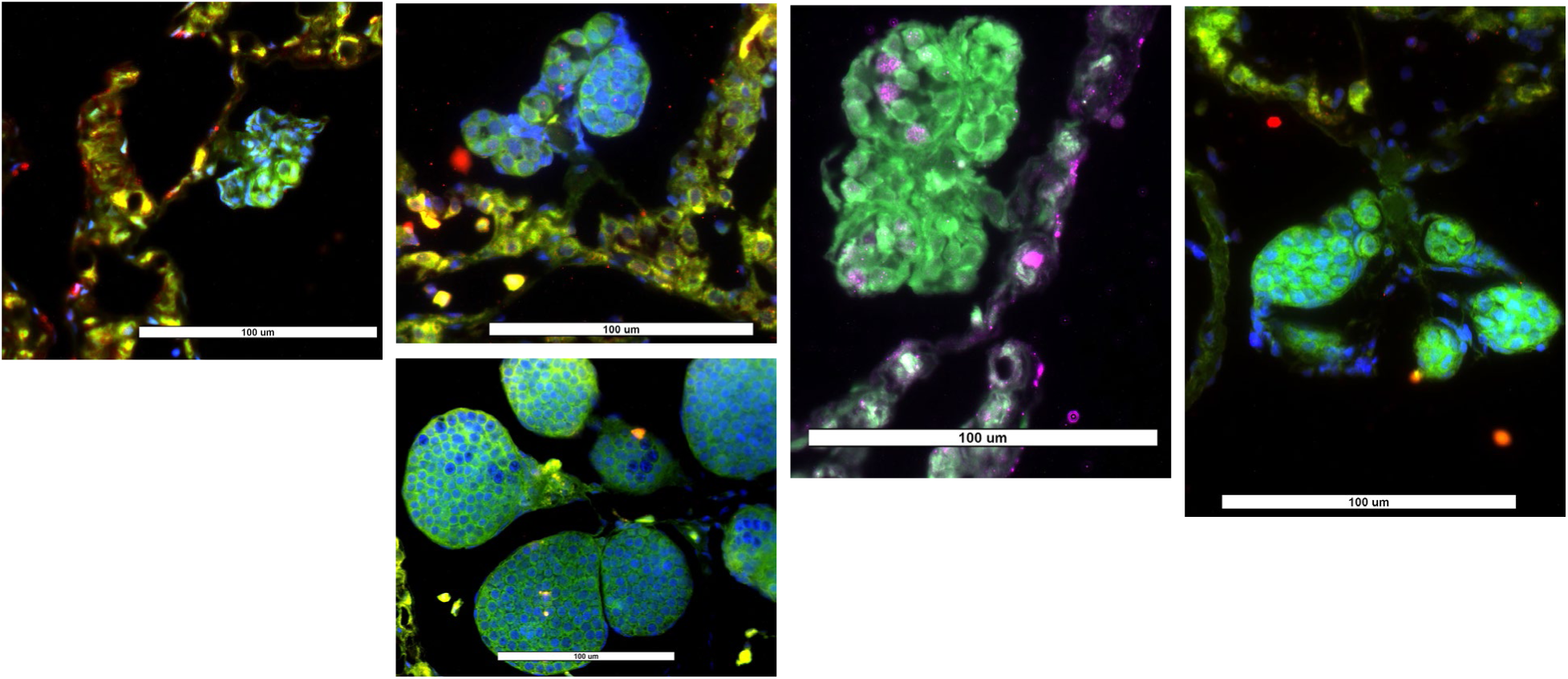
PIWIL1-immunoreactive (red) migrating primordial germ cells and spermatogonia in larval sea lamprey gonad. Blue: DAPI; Green: autofluorescence; Magenta: DAPI + PIWIL1; White: DAPI + PIWIL1 + autofluorescence; Orange and Yellow: PIWIL1 + autofluorescence.

**Figure 4.**
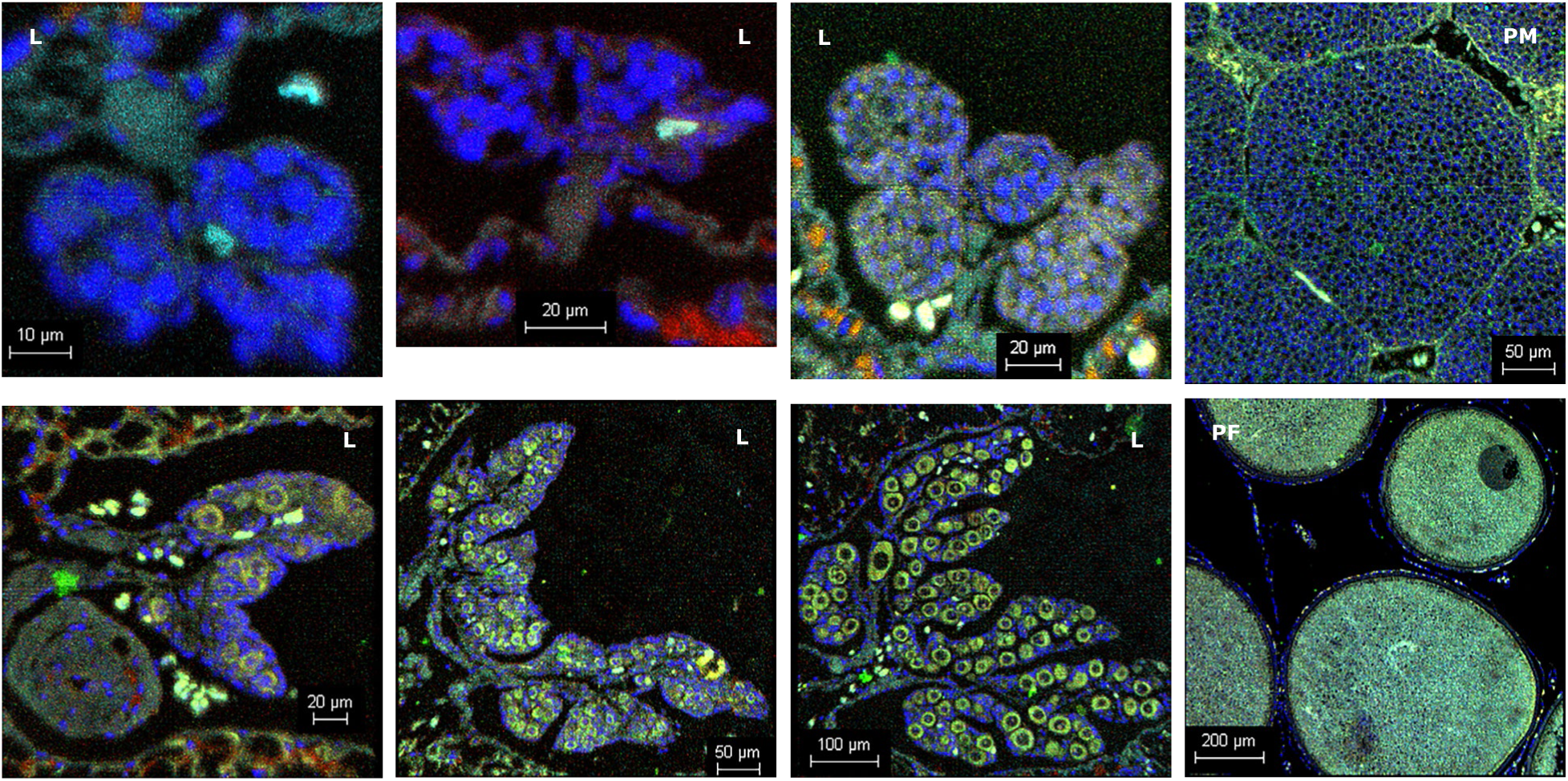
Images of gonadal tissues stain with HCR probes for BOLL (light blue), GLG1 (green), Piwil1 (yellow), and TEX11 (red). Blue: DAPI. L: larva; PF: preovulatory female; PM: prespermiating male.

**Figure 5.**
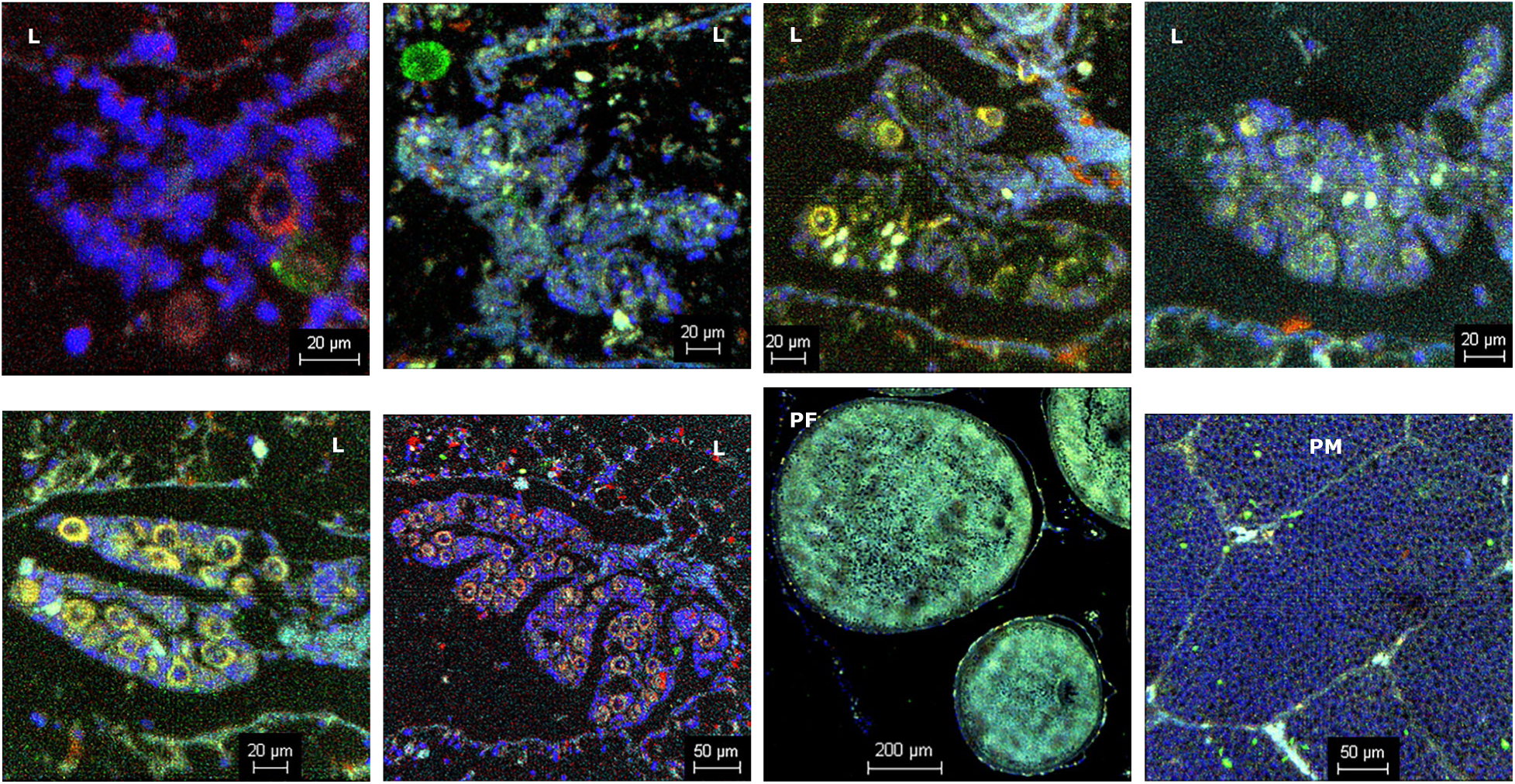
Images of gonadal tissues stain with HCR probes for CD9 (light blue), GLG1 (green), Piwil1 (yellow), and HSPA5 (red). Blue: DAPI. L: larva; PF: preovulatory female; PM: prespermiating male.

**Figure 6.**
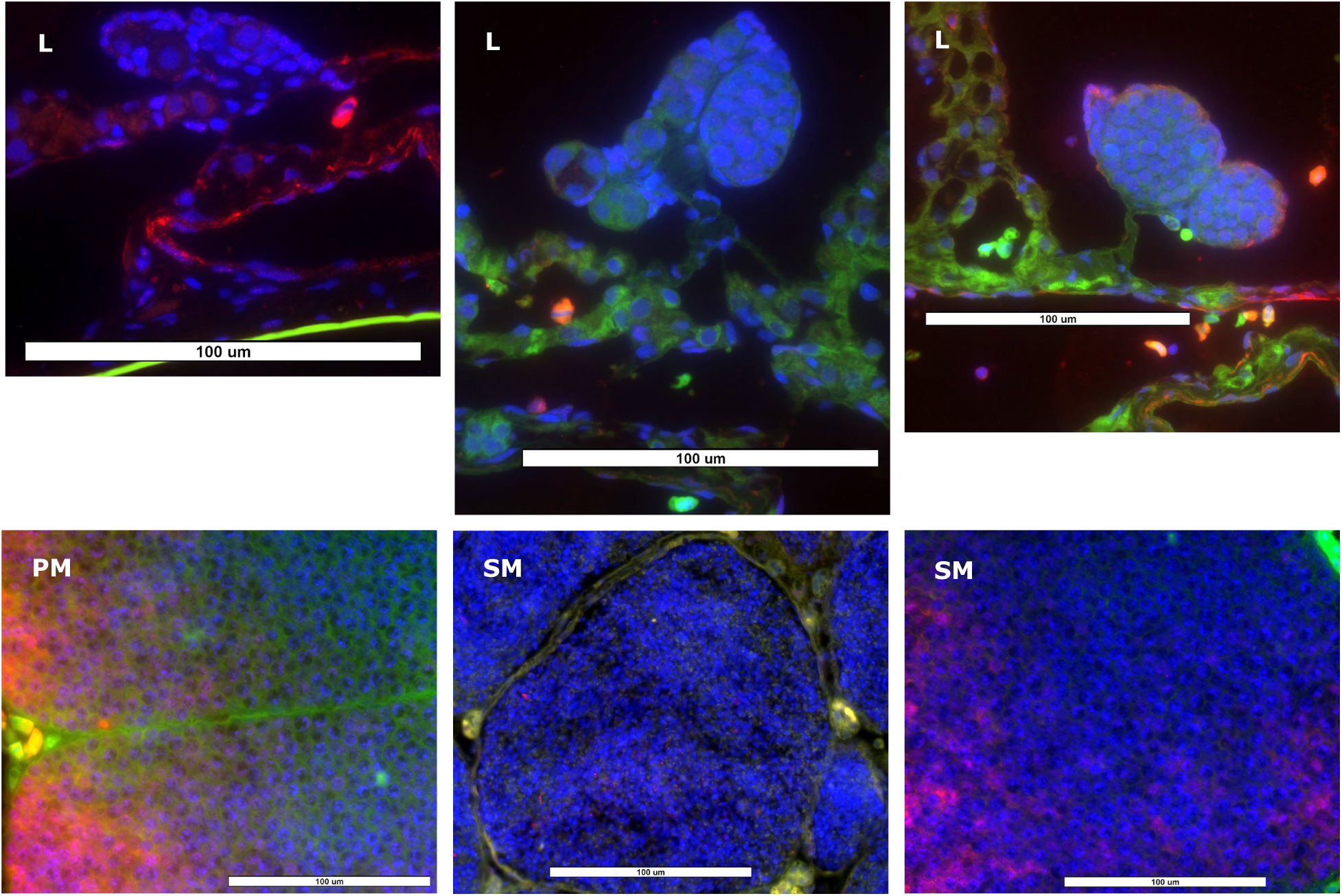
SYCP3-immunoreactive (red) migrating male germ cells and spermatocytes in sea lamprey gonad. Blue: DAPI; Green: autofluorescence; Magenta: DAPI + SYCP3; White: DAPI + SYCP3 + autofluorescence; Orange and Yellow: SYCP3 + autofluorescence.

### Pseudotime analysis identifies divergent male and female germ cell trajectories

We then isolated data from germ cells (clusters 5: oogonia/oocytes; 7: migrating primordial germ cells; 16: cytokinetic spermatocytes; 17: migrating male germ cells; 18: apoptotic spermatocytes; 19: GPCR-activated spermatocytes; Figure 1) to perform pseudotime trajectory analysis. These cells were re-clustered into 15 subclusters, and re-annotated as described above (Supplementary Tables S1, S12 and S13).

Genes with expressions changing along the developmental paths from primordial germ cells (PGCs) to mature germ cells were identified using pseudotime trajectory analysis (Figure 7, Supplementary Table S14). The top 50 genes exhibiting dynamic expressions along the male pathway (ending in SPC.cytokinetic cells) and female pathway (ending in ooctyes) are shown in Figures 8 and 9, respectively. GO analysis of gene expression changes along the male pathway were related to ribosomal function (GO: 0000027 ribosomal large subunit assembly; GO: 0000028 ribosomal small subunit assembly; GO:0000467 exonucleolytic trimming to generate mature 3’-end of 5.8S rRNA from tricistronic rRNA transcript), and translation (GO: 0002181 cytoplasmic translation; GO: 0006412 translation) (Supplementary Table S15). Genes involved in female pathway indicated significant increase in binding of sperm to zona pellucida (GO:0007339), egg coat formation (GO:0035803), prevention of polyspermy (GO: 0060468), and positive regulation of acrosome reaction (GO: 2000344) (Supplementary Table S15). In addition, we examined the genes differentially expressed between the primordial germ cells and germ cell cyst (oogonia stem cells). These genes were mainly involved in regulation of neurogenesis (GO: 0050767), cilium movement (GO: 0003341), and cilium assembly (GO: 0060271) (Supplementary Table S15).

**Figure 7.**
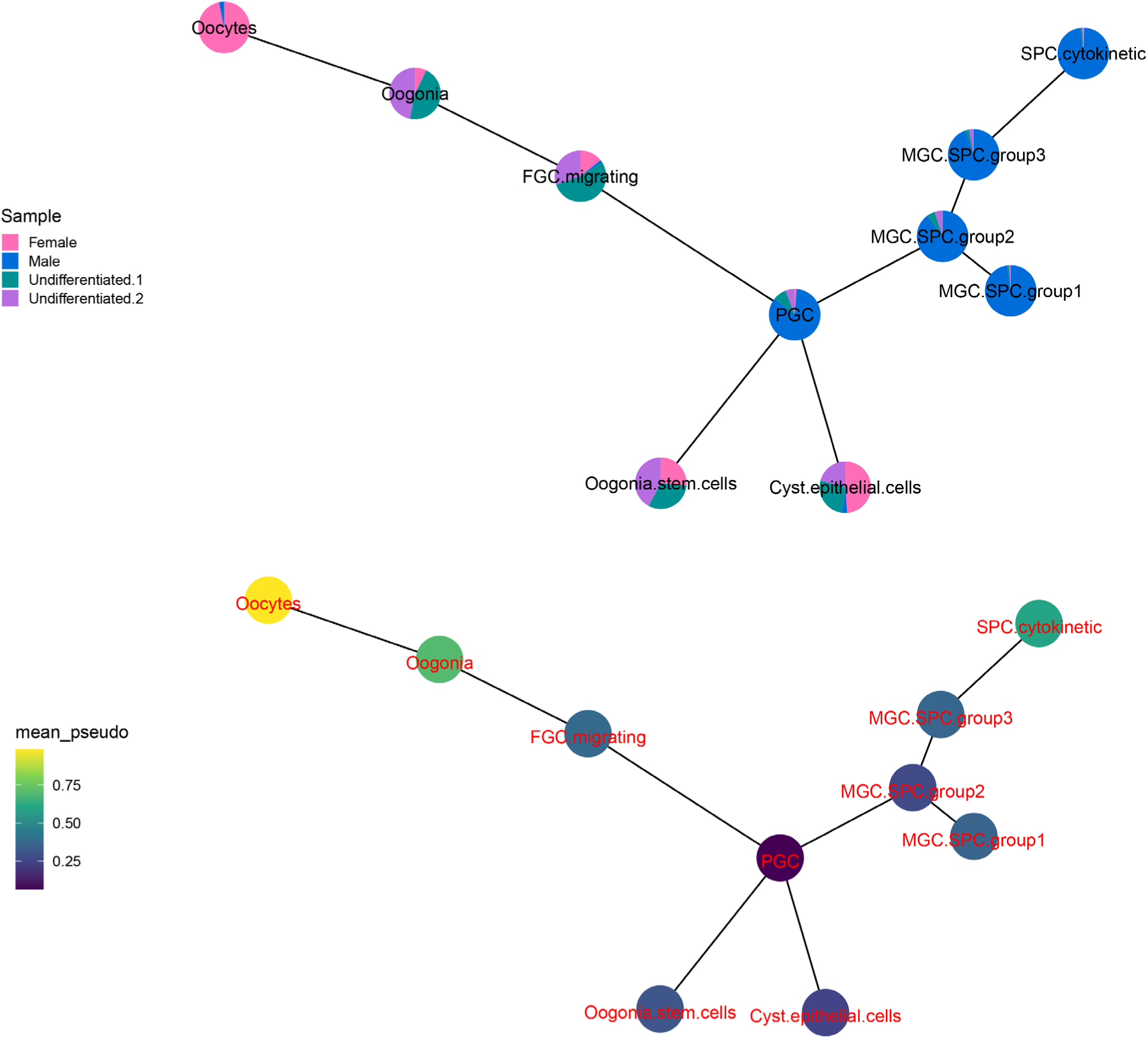
Pseudotime trajectory plotted with each cell cluster as a node. FGC: female germ cells; MGC: male germ cells; PGC: primordial germ cells; SPC: spermatocytes.

**Figure 8.**
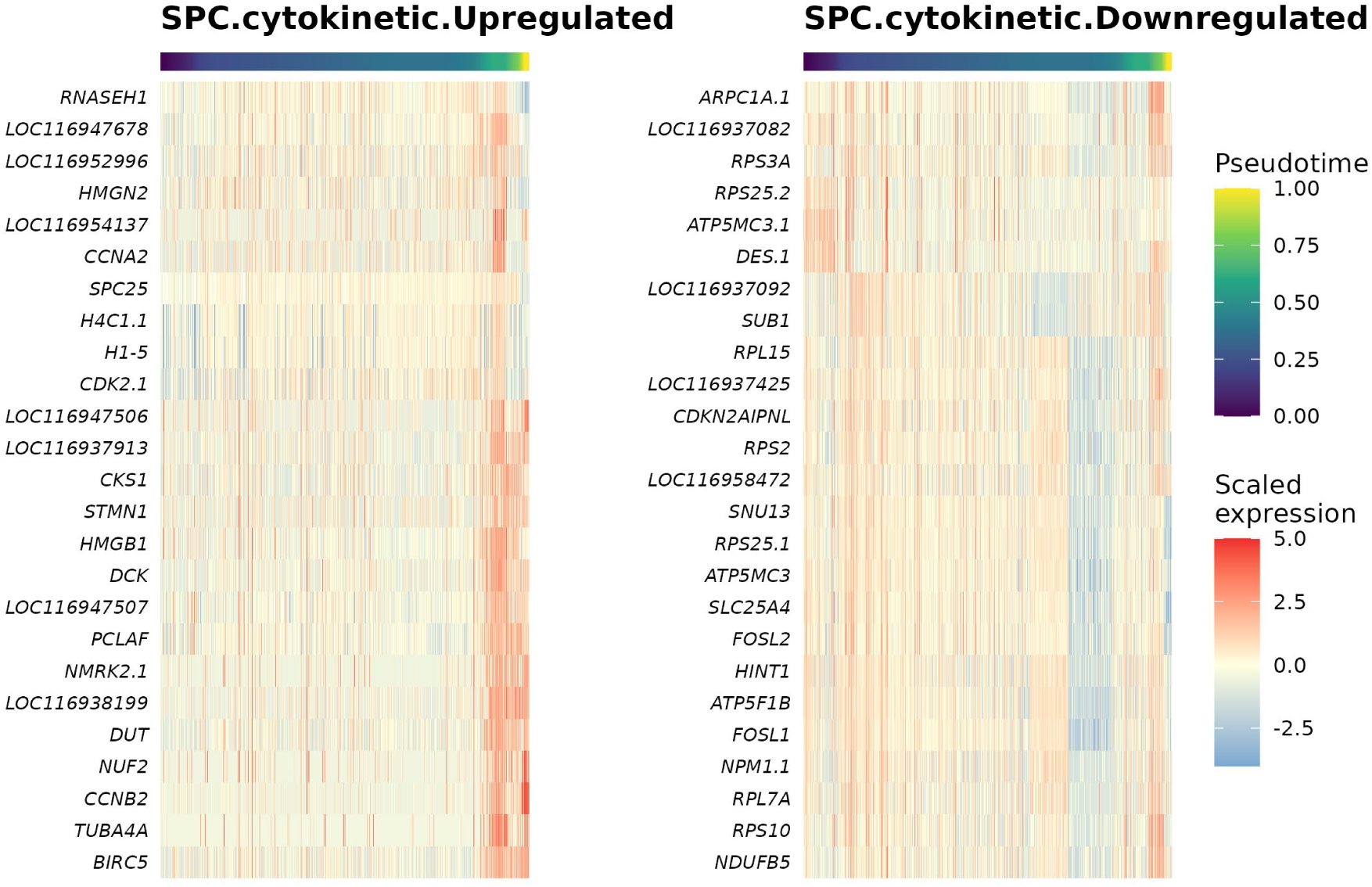
Heat maps of top 25 up- or down-regulated genes expressed in the pseudotime trajectory in the path from primordial germ cells to cytokinetic spermatocytes, i.e., male differentiation pathway. SPC: spermatocytes.

**Figure 9.**
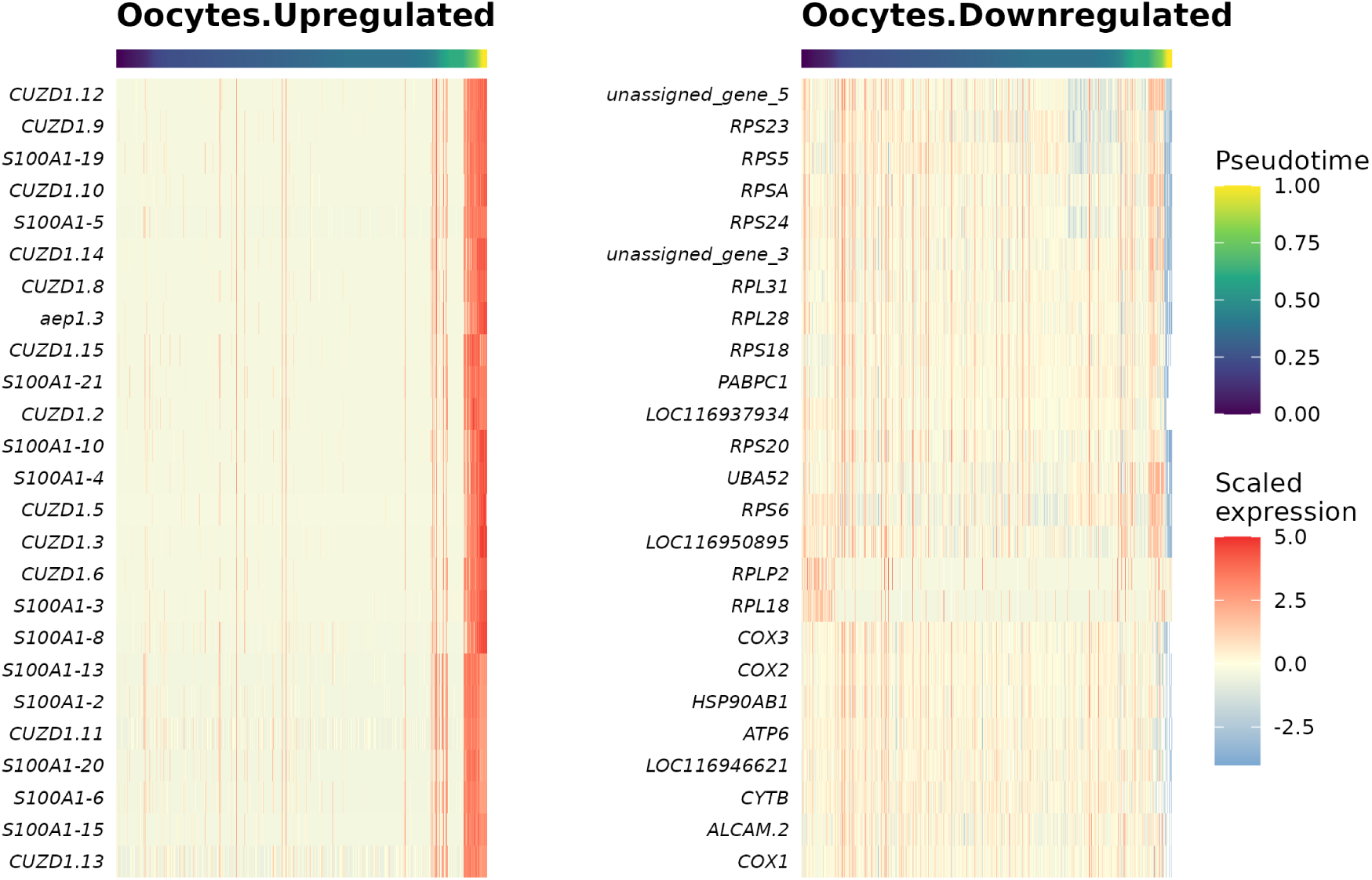
Heat maps of top 25 up-or down-regulated genes expressed in the pseudotime trajectory in the path from primordial germ cells to oocytes, i.e., female differentiation pathway.

In the initial cell clusters, WNT5 transcript variants showed cell type-specific expression patterns (Supplementary Figure S16). WNT5B (LOC116937338) was enrich in female germ cells (cluster 5: oogonia/oocytes), WNT5B.3 (LOC116949348) was enriched in somatic cells (cluster 2: mesonephric cells, cluster 3: migrating granulosa cells, and cluster 9: migrating pre-granulosa cells), and WNT5A-1 (LOC116937873) and WNT5B.1 (LOC116938414) were enriched in both female and male germ cells (Supplementary Figure S16). In the reclustered germ cell clusters, three WNT5 transcripts showed transcript-specific expression patterns (Figure 10). WNT5B was still enriched in migrating female germ cells (FGCs) and oogonia, whereas WNT5B.1 and WNT5A-1 were enriched in all male germ cells and a subset of migrating FGCs and oogonia (Figure 10). These results indicate transcript-specific, sexually dimorphic WNT5 expression in the sea lamprey gonad during sexual differentiation, with one transcript restricted to the female lineage and two shared between the female and the male germ cell lineage. Designing transcript-specific probes was not feasible because the sequences are highly similar among these transcripts. Consistent with this expression across both lineages, WNT5B immunoreactivity was detected in migrating cells, within the germ cell cyst, and in both spermatocytes and oocytes (Figure 11). Moreover, WNT5 transcripts have distinct expression over pseudotime (Figure 12). Consistent with their cell-type expression patterns, WNT5B.1 and WNT5A-1 increased along the female path until the oogonia and then decreased in expression in more mature oocytes, whereas along the male path these genes peaked at an earlier pseudotime (Figure 12). WNT5B similarly peaked along the female path before decreasing in oocytes, while its expression remained unchanged on the male path (Figure 12). While both have relatively low expression, WNT5B.2 (LOC116947519) and WNT5B.3 peaked early in pseudotime along the female path with higher expression in migrating FGCs and oogonia stem cells (Figure 12). Along the male path WNT5B.3 decreased with pseudotime while WNT5B.2 expression remained flat (Figure 12).

**Figure 10.**
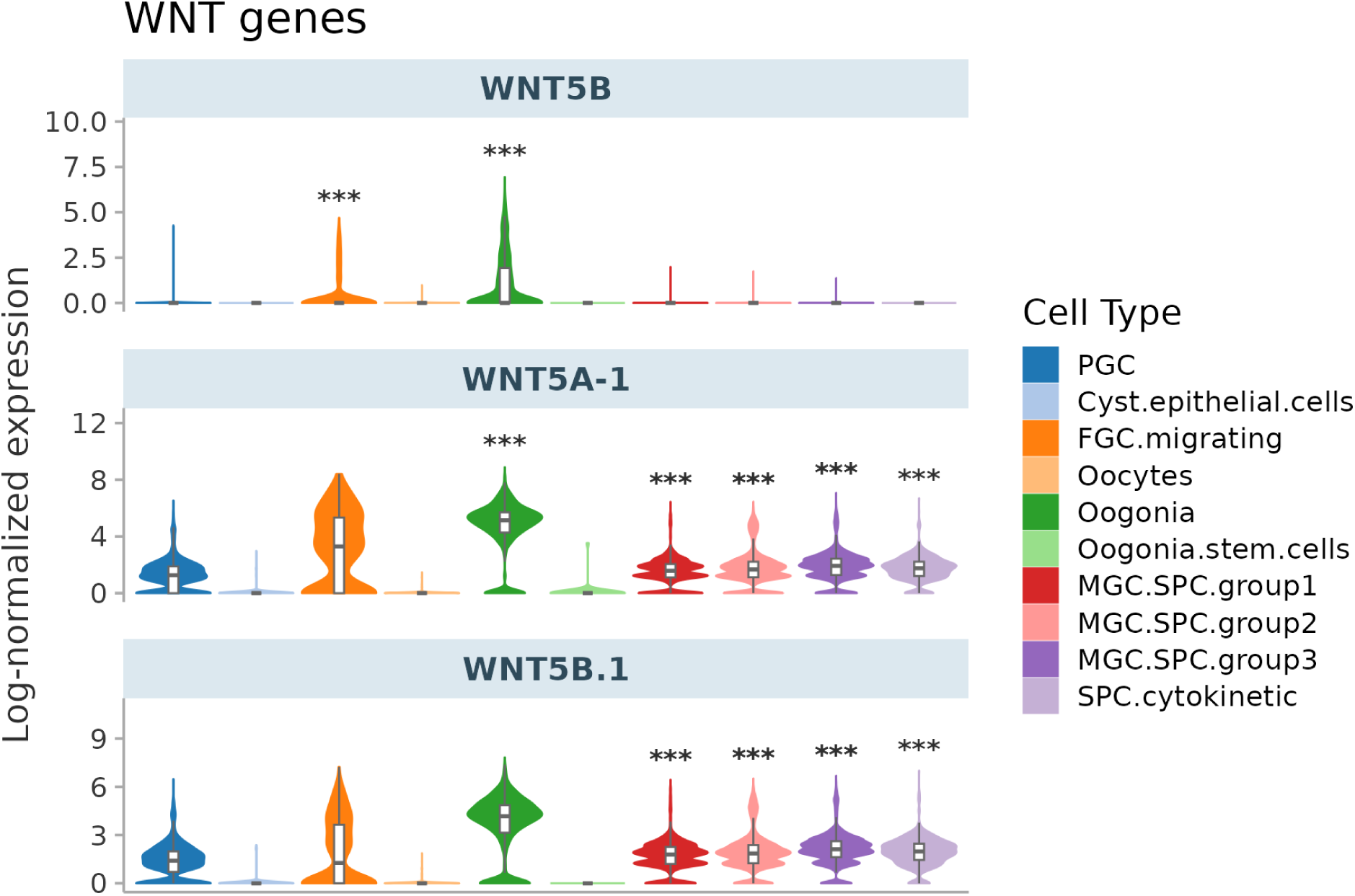
Violin plots of WNT5A and WNT5B gene expressions in germ cell clusters. Note that WNT5B transcript show sexually dimorphic expression pattern (mainly in female germ cells).

**Figure 11.**
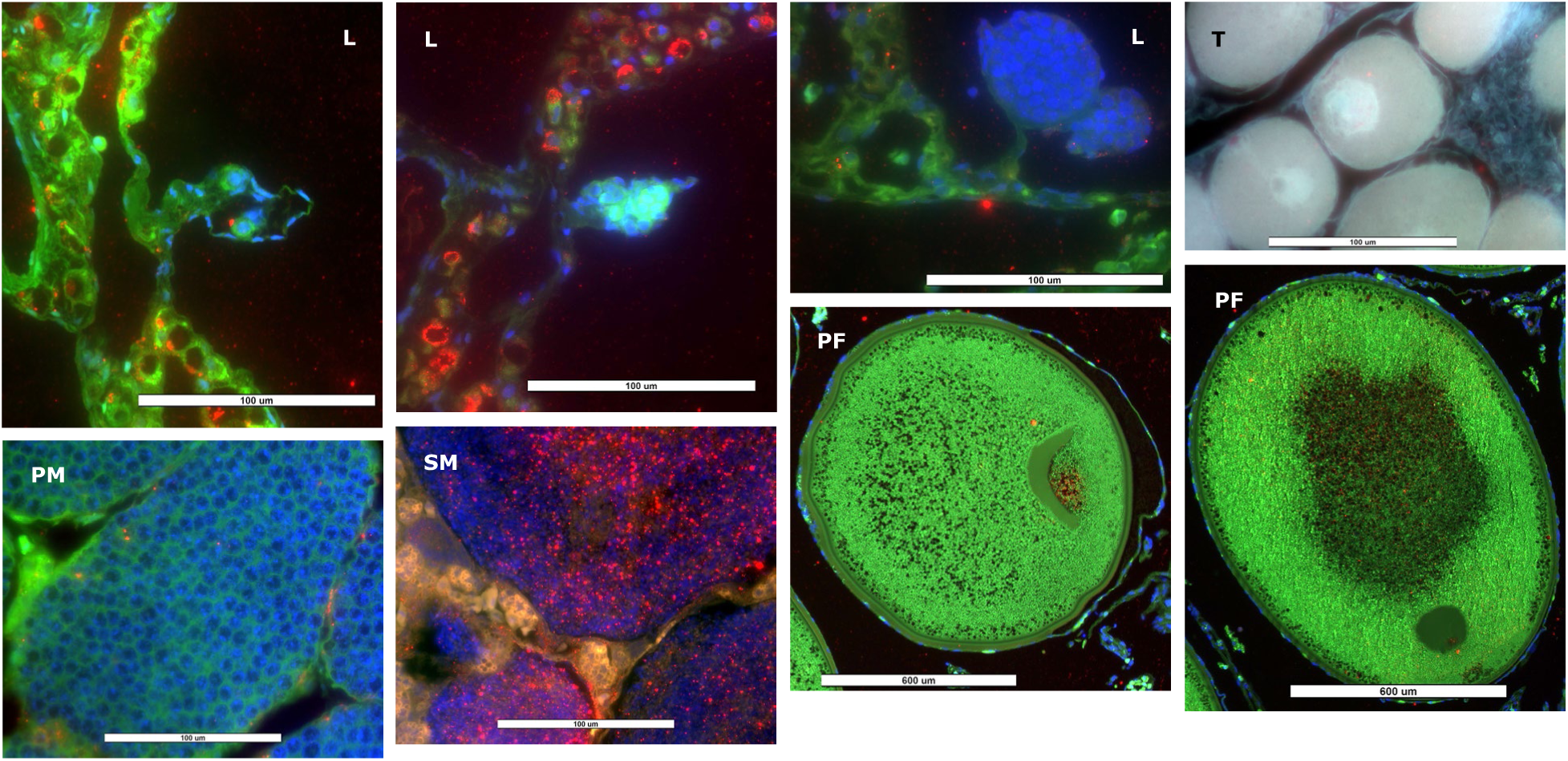
WNT5B-immunoreactive (red) germ cells in sea lamprey gonad at different developmental stage. Blue: DAPI; Green: autofluorescence; Magenta: DAPI + WNT5B; White: DAPI + WNT5B + autofluorescence; Orange and Yellow: WNT5B + autofluorescence.

**Figure 12.**
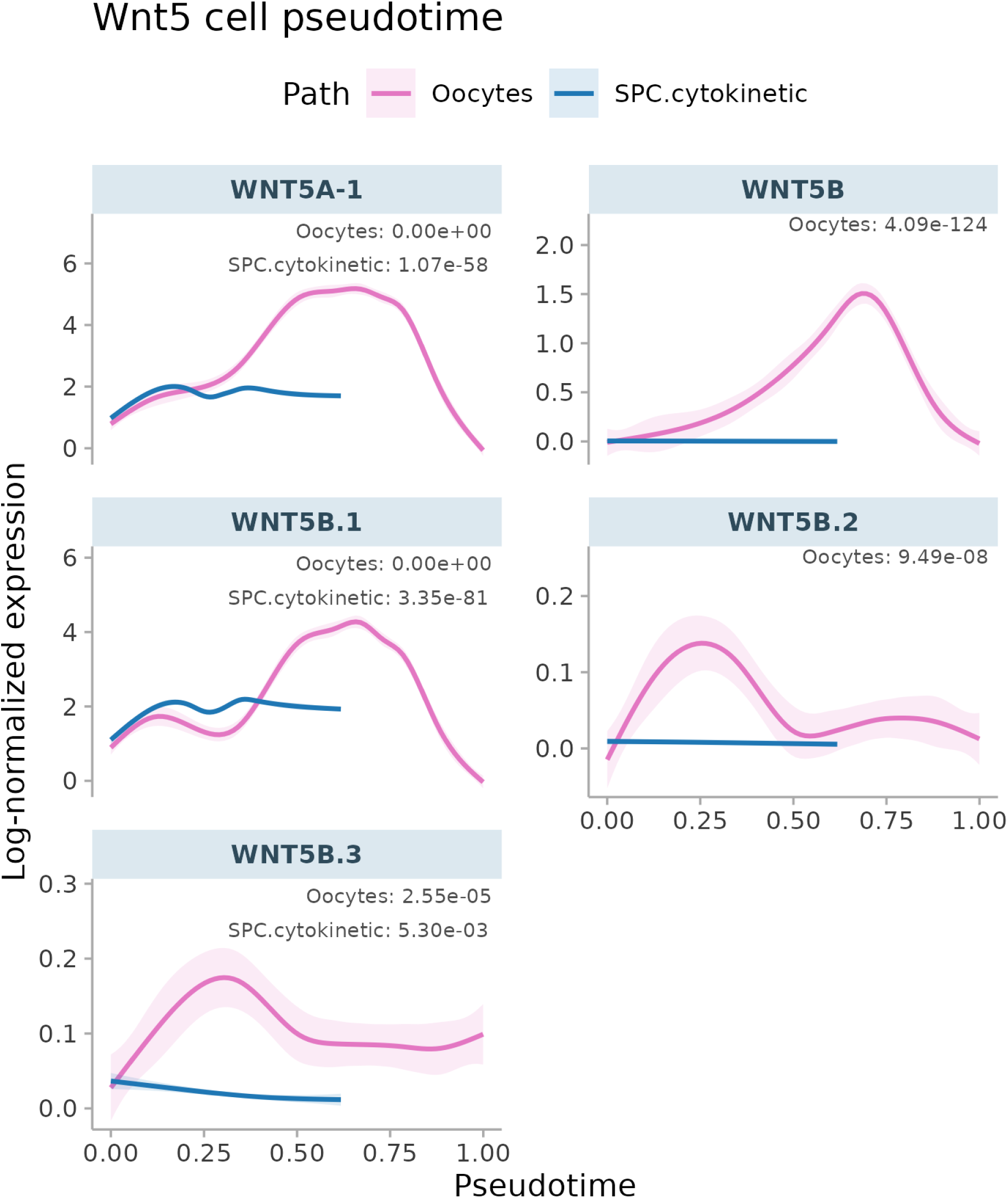
Pseudotime expression patterns of WNT5A and WNT5B transcripts in male and female differentiation pathways.

Given these results, our working hypothesis is that WNT signaling pathways, which can be modulated by environmental factors like iron concentration (Baschant et al., 2016), play important roles during germ cell development (Figure 13), including during cell fate commitment. However, whether these genes or signaling pathways play a role in sea lamprey sex determination or sexual differentiation requires further investigation.

**Figure 13.**
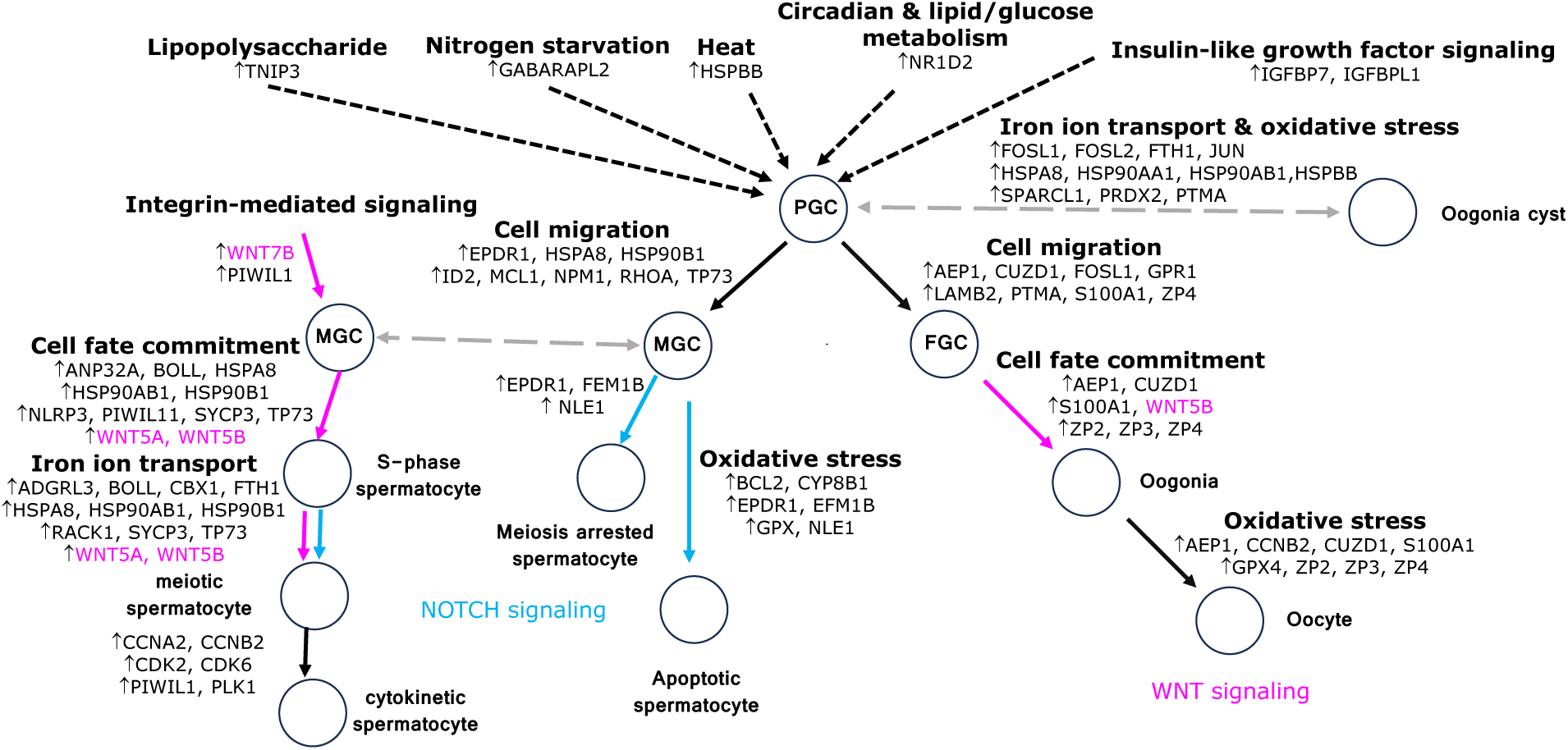
A diagraph showing a working hypothesis for sea lamprey germ cell development. FGC: female germ cells; MGC: male germ cells; PGC: primordial germ cells.

### Germline-specific genes are expressed in germ cells

Current sea lamprey genome contains one germline-specific chromosome (Chromosome 81; Timoshevskaya et al., 2023). Of the 13 genes located on chromosome 81with detected expression in our PIP-seq experiment, 11 showed enrichments in germ cell clusters. For example, HYKK (LOC116957691), HYKK.5 (LOC116957686) and HYKK.6 (LOC116957689) were enriched in both male and female germ cells, while HYKK.4 (LOC116957684) is only enriched in oogonia (Figure 14). Other chromosome 81 genes such as SPOPL (LOC116957694), CDH4.3 (LOC116957687), SPOPL.1 (LOC116957693), and SPOPL.2, are also enriched in germ cells of both sexes (Figure 14).

**Figure 14.**
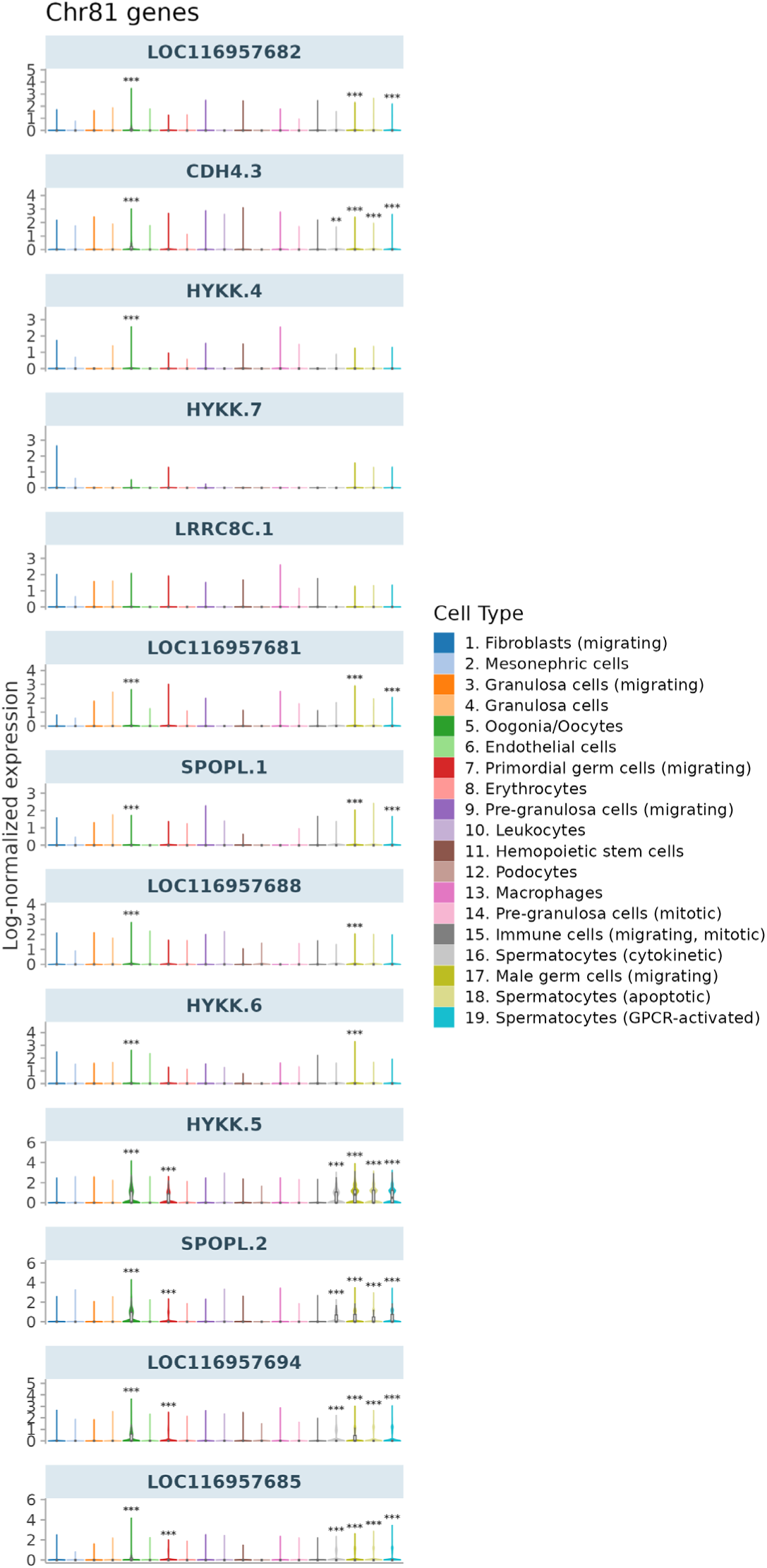
Expression patterns of genes located on the germline-specific chromosome (Chromosome 81) in all cell clusters.

A bulk RNA-seq study of gonads from larval chestnut (*Ichthyomyzon castaneus*) and northern brook lamprey (*I. fossor*) undergoing mitotic proliferation or the early stages of meiosis I (Ajmani et al., 2021), showed ZP2, ZP4, and NXPH1 expressions in differentiated female gonads. We found the same genes enriched in oocytes/oogonia in our PIP-seq results. Moreover, Ajmani et al. showed SPAG16 expressions in differentiated male gonads. Indeed, the highest median expression of SPAG16 is observed in male germ cells in our PIP-seq results. Furthermore, we found IL8, PFN2, and STC2 enriched in migrating PGC; IL8, PFN2 (LOC116952631, LOC116957159), RPA1 (LOC116939309, LOC116950263), MMP19, IGF1, and IGF1R enriched in somatic cells; and ZP, IGFBP5, IQUB, IQCG, MNS1 and ALOX5 enriched in somatic and germ cells (Supplementary Figure S17). Our PIP-seq results are consistent with bulk RNA-seq results, and further distinguished genes expressed in germ cells vs. somatic cells.

### GnRH, growth factors, and their receptors may be important for germ cell development in sea lamprey

In another bulk RNA-seq study of germline-specific genes in sea lamprey genome, GO terms of these genes were associated with WNT signaling pathway, insulin/insulin growth factor (IGF) pathway, gonadotropin-releasing hormone receptor (GnRHR) pathway, and transforming growth factor-beta (TGFβ) signaling pathway (Yasmin et al., 2022). We also found that IGF, GnRH and their receptors were expressed in germ cells and surrounding somatic cells in sea lamprey gonad, suggesting potential interactions between germ cells and gonadal milieu.

GnRH2.1 expression was enriched in germ cells, whereas GnRH receptors and GnRH2 were expressed in somatic and germ cells (Supplementary Figure S18). Insulin growth factor 1-1 (IGF1-1) and IGF1R expressions were enriched in germ cells and somatic cells (Supplementary Figure S18). However, Insulin-like growth factor 1 (IGF1), IGF1R.1, and IGF1R.2 were differentially expressed in somatic cells (Supplementary Figure S18). In vertebrates, IGF1R is expressed during the first phase of oocyte growth, and expression level is directly related to number of oocytes per section (Spice et al., 2014). IGF1 is associated with increased growth and fecundity (but reduced lifespan), and it also stimulates the production of sex steroids (Dantzer & Swanson, 2012). Successful execution of the germ cell developmental program requires ongoing juxtacrine, paracrine, and endocrine signaling between germ cells, supporting somatic cells, and the pituitary gland (DeFalco et al., 2015; Franca et al., 1998; Grisword, 1995; Meng et al., 2000; O’Shaughnessy et al., 2008; Sharpe, 1986). We observed an enrichment of GnRH and GnRHR in pre-granulosa and germ cells, suggesting a possibility that locally produced GnRH may regulate early germ cell development and may not require pituitary GnRH inputs. Further analysis is required to confirm their involvement during sea lamprey sexual differentiation.

TGFβ superfamily signaling is involved in different reproductive processes, such as sex determination (Rajendiran et al., 2021), sexual differentiation (Kwan & Patil, 2019), gametogenesis (Rajendiran et al., 2021), fertilization and sexual behavior (Padilla et al., 2021). TGFβ and bone morphogenetic protein (BMP) signaling pathways play essential roles in regulating embryonic and adult stem cell niches. In zebrafish, follicle-stimulating hormone regulates different members of the TGFβ superfamily, which play important roles in the balance between spermatogonia proliferation and differentiation in the male germ stem cell niche (da Costa et al., 2024). We found endothelial, pre-granulosa and granulosa cells differentially expressed TGFβ receptor II (TGFβR2) and its ligand (TGFβ2) (Supplementary Figure S19). These cells may be able to self-regulate the effects of TGFβ signaling. Moreover, oogonia/oocytes also differentially expressed downstream SMAD6 and SNIP1 (inhibitory modulators), whereas pre-granulosa cells and male germ cells differentially expressed SMAD3 (Supplementary Figure S19). BMPR2 expression was enriched in germ cells, mesonephric, pre-granulosa, and granulosa cells (Supplementary Figure S19). Interestingly, mesonephric and germ cells also differentially expressed KCP that positively regulates BMP signaling and BMPER that negatively regulate BMP functions, as well as BMP2K that inhibits differentiation (Supplementary Figure S19). Oogonia/oocytes also differentially express BRINP3 that inhibits cell cycle transition, promotes cell proliferation, migration and invasion (Supplementary Figure S19). On the contrary, pre-granulosa and granulosa cells, and male germ cells differentially expressed BRINP1 that inhibits cell cycle progression and G1/S transition (Supplementary Figure S19). A deeper understanding of TGFβ family-mediated intercellular interactions within the sea lamprey gonad will require characterization of the spatiotemporal signaling networks that coordinate communication between germ cells and supporting somatic cells.

### Sex steroids and bile acids may be important for sexual differentiation in sea lamprey

Sex steroids play important roles in sexual differentiation in most vertebrates. Steroidogenic enzymes such as aromatase CYP19A1, 3β-hydroxysteroid dehydrogenase (HSD3b1), and 11-hydroxylase (CYP11b2.1) appear to play important roles during the critical period of sex determination and sexual differentiation in many species (Docker et al., 2019). However, Docker was unable to direct the course of gonadal differentiation by injecting lamprey with gonadal steroids. Interestingly, we found that mesonephric and germ cells differentially expressed estrogen-regulated ESRRG, ESRRG-1, and ESRRG-2, whereas pre-granulosa and granulosa cells differentially expressed CYP17A1 and androgen receptor NR5A1 (Supplementary Figure S20). While the course of sexual differentiation may not be susceptible to manipulation by exogenous steroid hormones, the influence of endogenous steroid hormones required further investigation (Docker et al., 2019).

### Sex determination and sexual differentiation in sea lamprey may be in a transitional state

Sex has a genetic basis among the lower vertebrates despite the frequent absence of recognizable sex chromosomes (Yarnamoto, 1953; Richards & Nace, 1978; Hunter et al., 1983; Yamazki, 1983; Nagahama et al., 2021). The underlying sex regulator genes appear to have converged on the doublesex and male abnormal-3 (Mab-3)-related transcription factor (Dmrt) gene family in many species (Kopp, 2012; Zarkower, 2001). Other transcription factors known to influence sexual differentiation in mammals include FOXL2 (forkhead box L2) in ovarian development, and SOX9 and steroidogenic factor (SF1) in testicular development (Bulun et al., 2003; Wilhelm et al., 2007; Sandra & Norma, 2010). In our PIP-seq data, FOXL2 was enriched in somatic cells such as fibroblasts, pre-granulosa and granulosa cells, which were mostly captured from females and larvae, and very few from males. SOX9 was enriched in fibroblasts and both male and female germ cells (Supplementary Figure S20). SF-1, the protein encoded by NR5A1 gene (Supplementary Figure S20), is enriched in pre-granulosa and granulosa cells. However, DMRT1 is not annotated in the sea lamprey genome, its expression could not be unequivocally assessed in the cells profiled in this study. Based on the available expression patterns, these genes may not exert similar regulatory effects on sexual differentiation in sea lamprey as in higher vertebrates.

Although the mechanisms underlying germ cell fate specification are diverse in animals, several properties of PGCs appear to be conserved. One property is that in PGCs the somatic differentiation program was repressed. In C. elegans and *D. melanogaster*, this is achieved through transient global repression of the RNA polymerase II (RNAPII) activity by PIE-1 and the polar granule component (Nakamura et al., 1996; Seydoux et al., 1996). PIE-1 inhibits the transcriptional initiation and elongation activity of RNAPII by distinct mechanisms (Ghosh & Seydoux, 2008). PIE-1 inhibits the positive transcriptional elongation factor b (P-TEFb), which promotes the transcriptional elongation activity of RNAPII by phosphorylating the Ser-2 of the carboxy-terminal domain (CTD) of RNAPII (Zhang et al., 2003). The polar granule component inhibits the recruitment of p-TEFb to transcriptional sites, therefore blocking the transcriptional initiation (Hanyu-Nakamura et al., 2008). In contrast, in mice, germ cell specification requires active transcription, and the transcriptional repressor BLIMP1 specifically shuts off gene expression for the somatic mesodermal program (Kurimoto et al., 2008). In sea lamprey, SRCAP transcripts (LOC116950250), with 67% identity with PIE-1 mRNA, were enriched in spermatogonia. CDK9 transcripts, with 70% identity with p-TEFb, were enriched in somatic and germ cells (Supplementary Figure S21). However, BLIMP1/PRDM1 (LOC116952900 and LOC116952967) were enriched in somatic and germ cells. These results suggest that the mechanism of PGC development in sea lamprey may share characteristics of both invertebrate and vertebrate modes of germ cell specification, potentially representing an intermediate evolutionary state.

## Conclusion/Summary

Sea lamprey sex determination and sexual differentiation likely involve many genes that interact with environmental factors, representing a transitional state between environmental sex determination and genetic sex determination. This study provides a resource for future studies of germ cell biology and may inform population control effort in sea lamprey, a notorious invasive species in the Great Lakes.

## Acknowledgements

We thank the Great Lakes Fishery Commission for funding support for Dr. Weiming Li. We thank Dr. Nicholas S. Johnson, Braden A. Idalski, and the staff of U.S. Geological Survey, Hammond Bay Biological Station, Great Lakes Science Center for lamprey procurement and sample collections. We thank the staff in the Flow Cytometry and Bioinformatics Core Facilities, and the Investigative Histopathology Laboratory of Michigan State University for their technical support.

