## Supplementary figures and images for "Gonadal PIP-seq reveals genes involved in germ cell development and sexual differentiation in sea lamprey (*Petromyzon marinus*)"

### Supplementary Figure S6

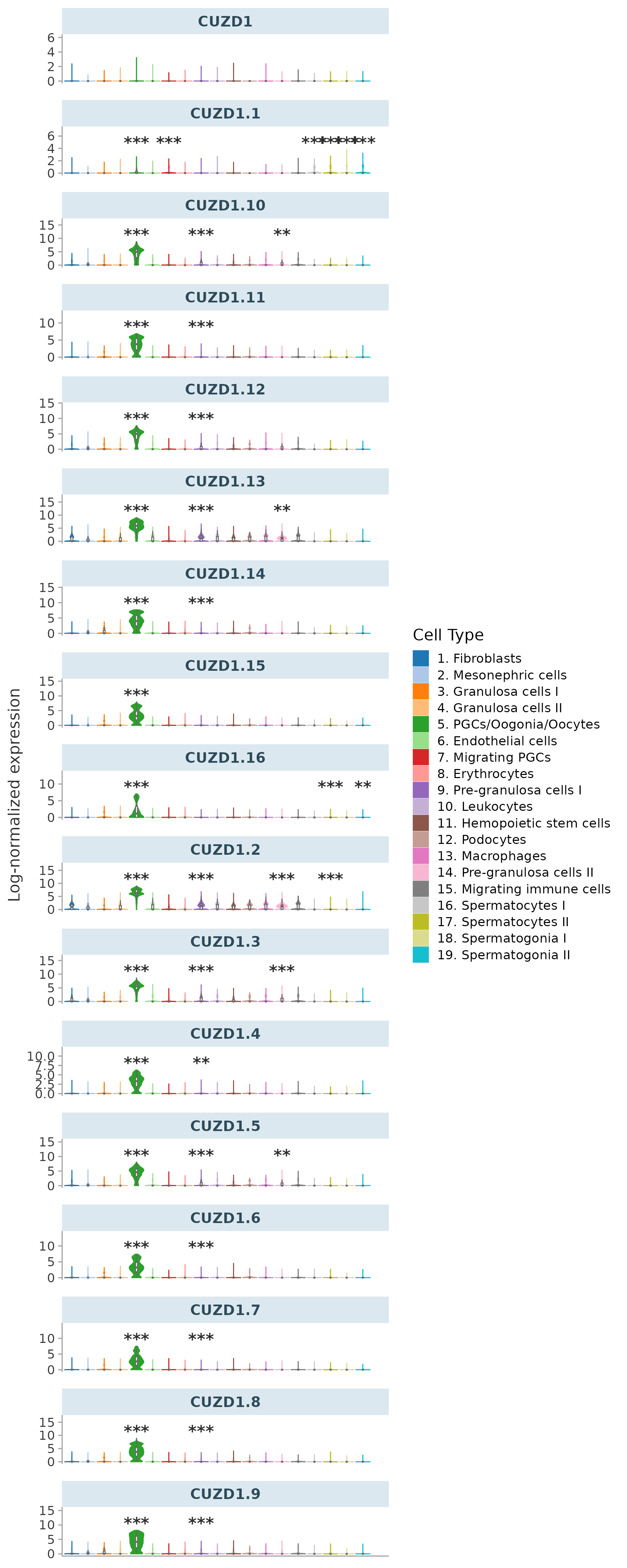

### Supplementary Figure S7

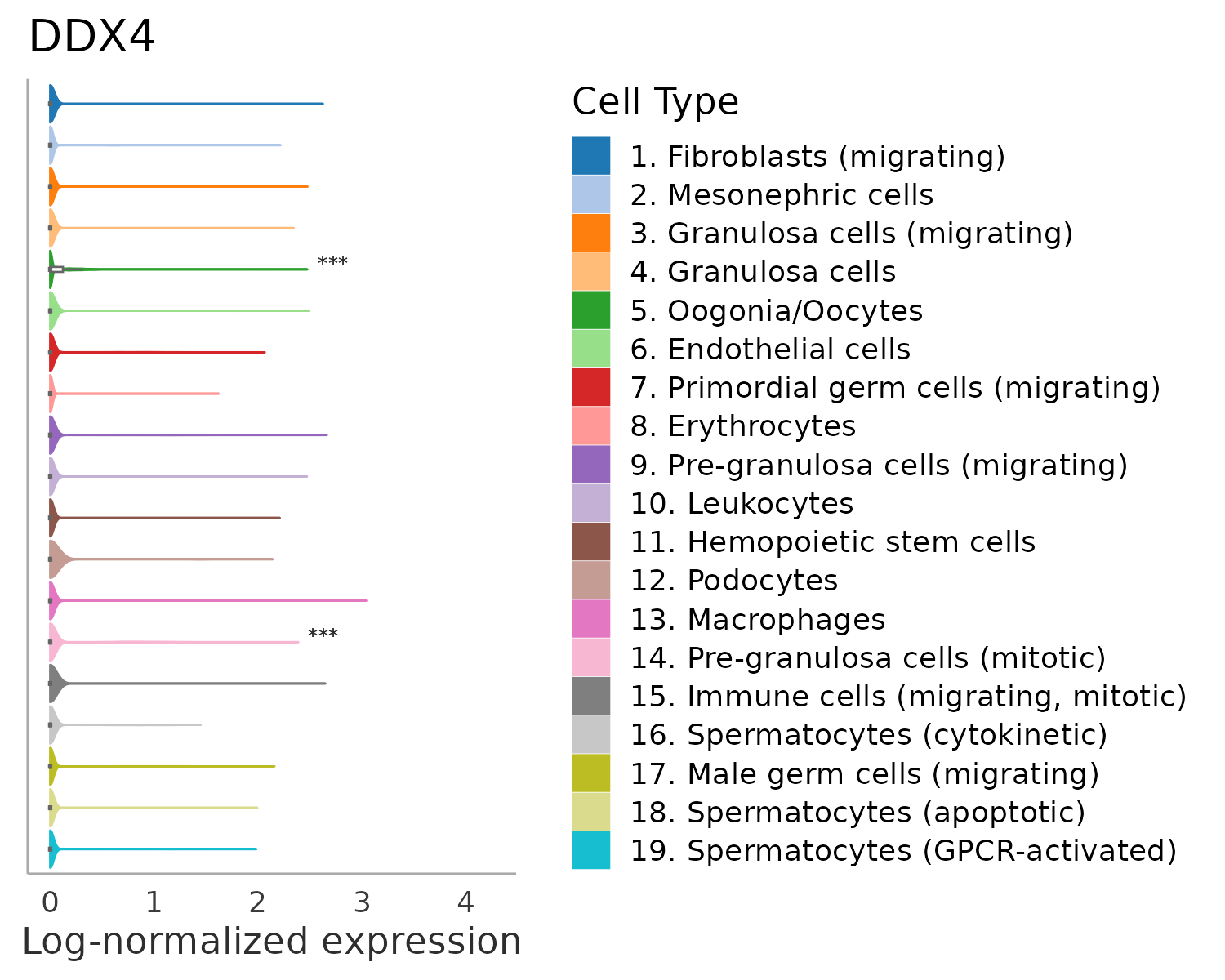

### Supplementary Figure S8

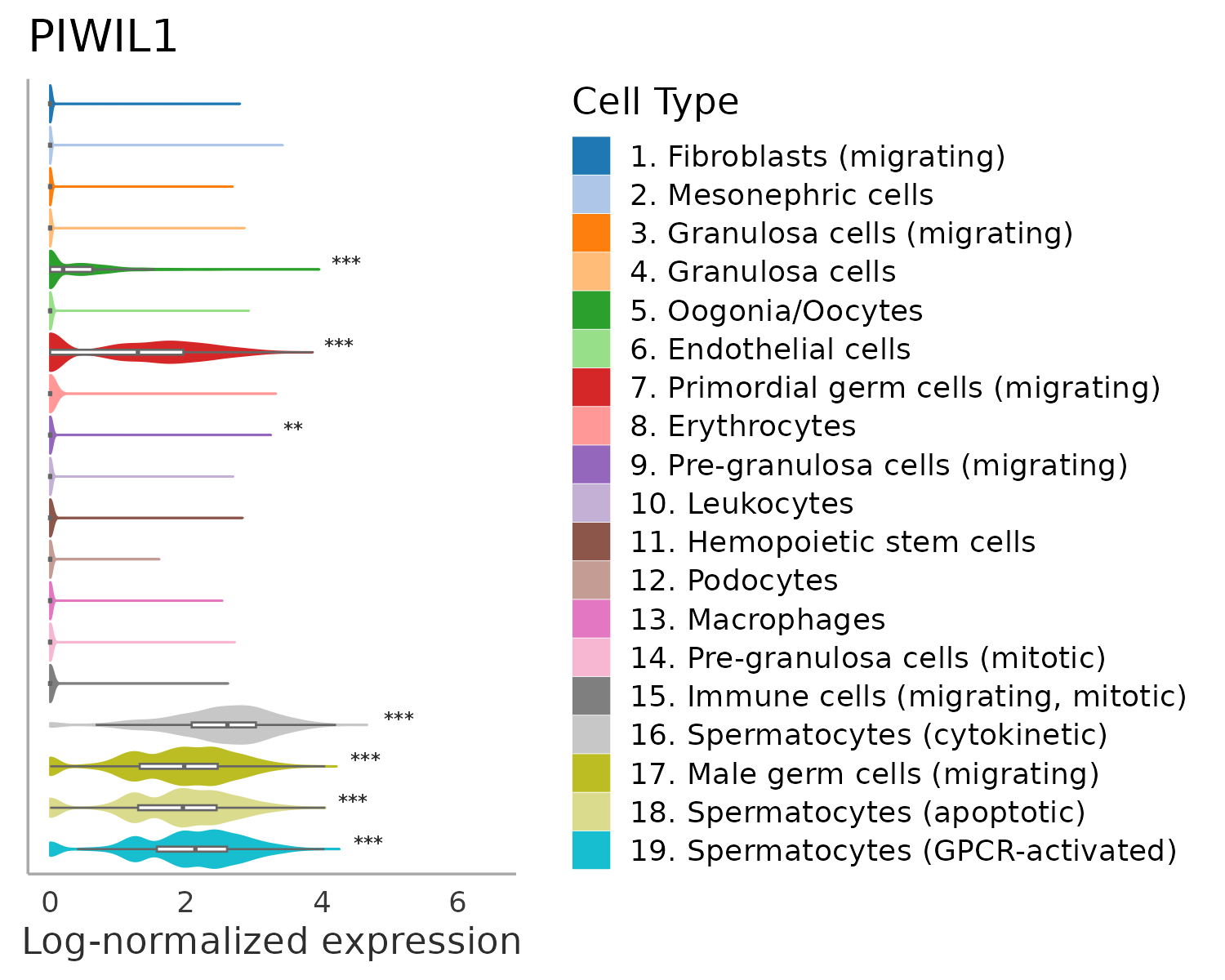

### Supplementary Figure S9

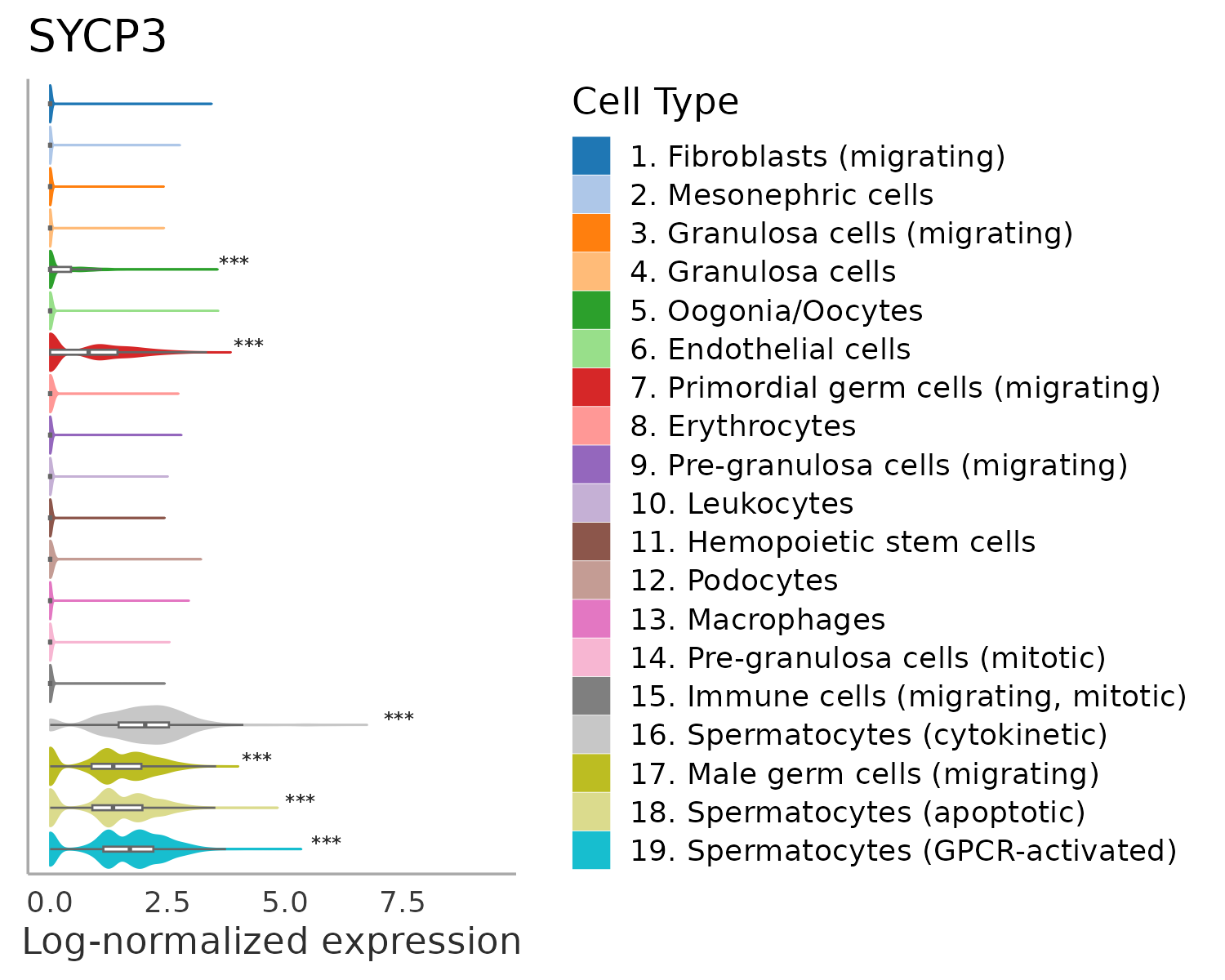

### Supplementary Figure S11

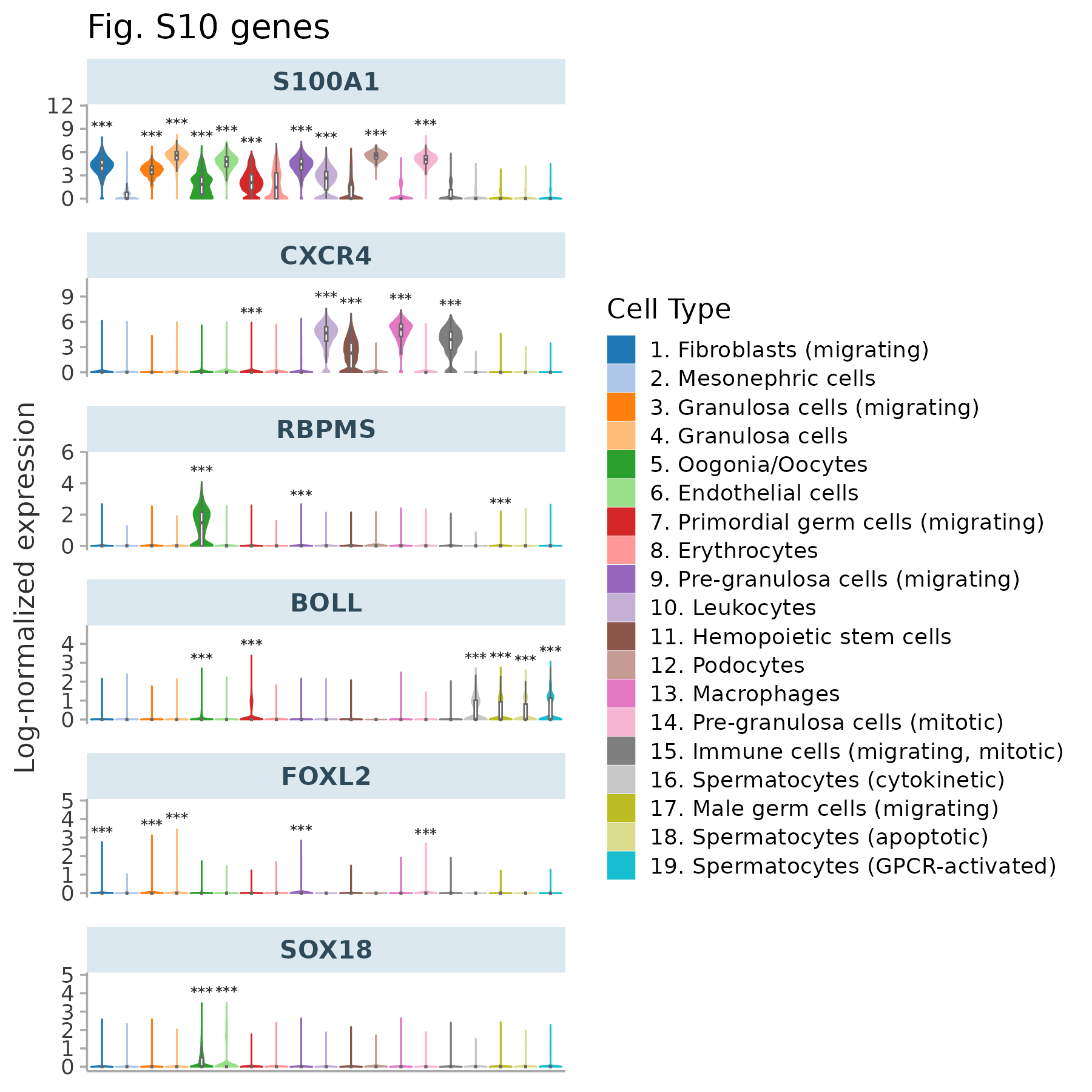

### Supplementary Figure S16

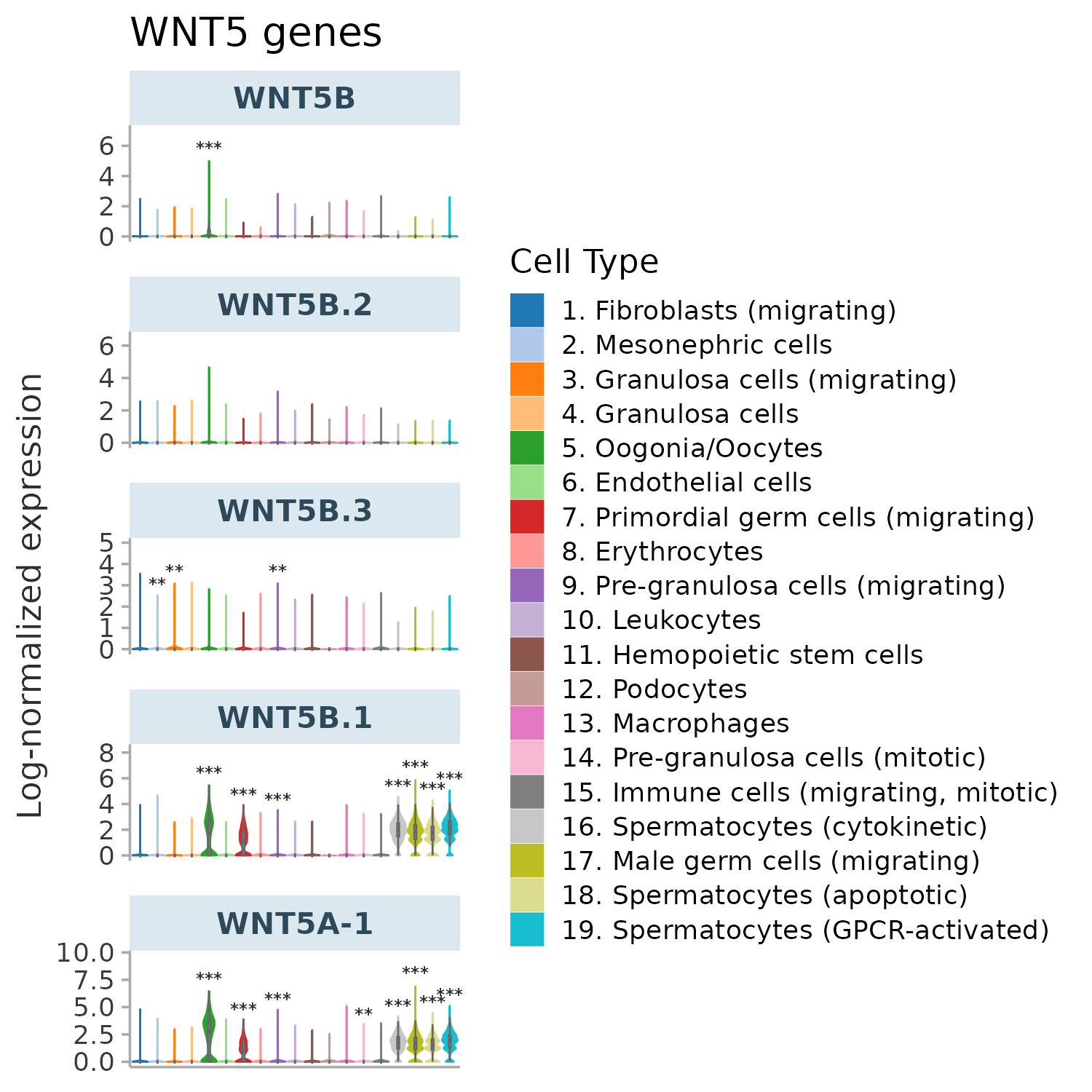

### Supplementary Figure S17

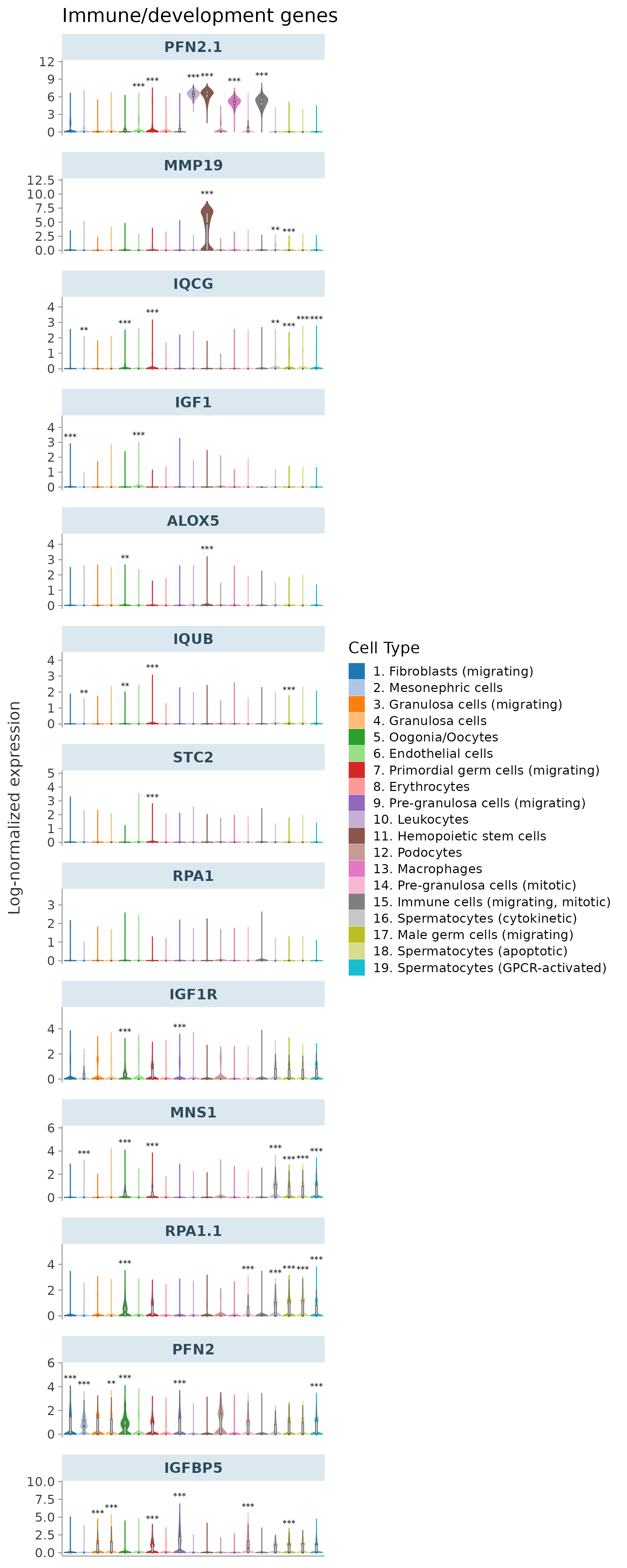

### Supplementary Figure S18

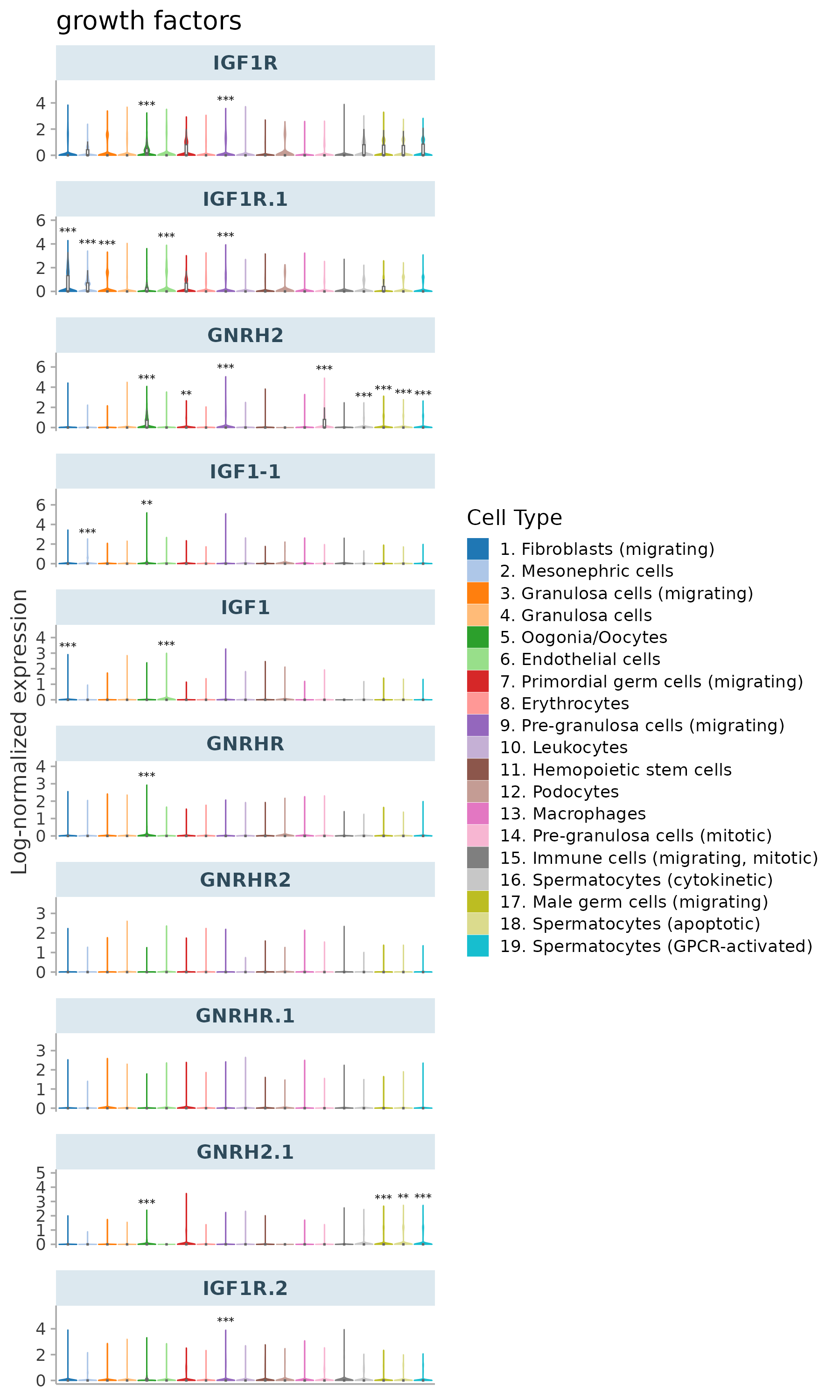

### Supplementary Figure S19

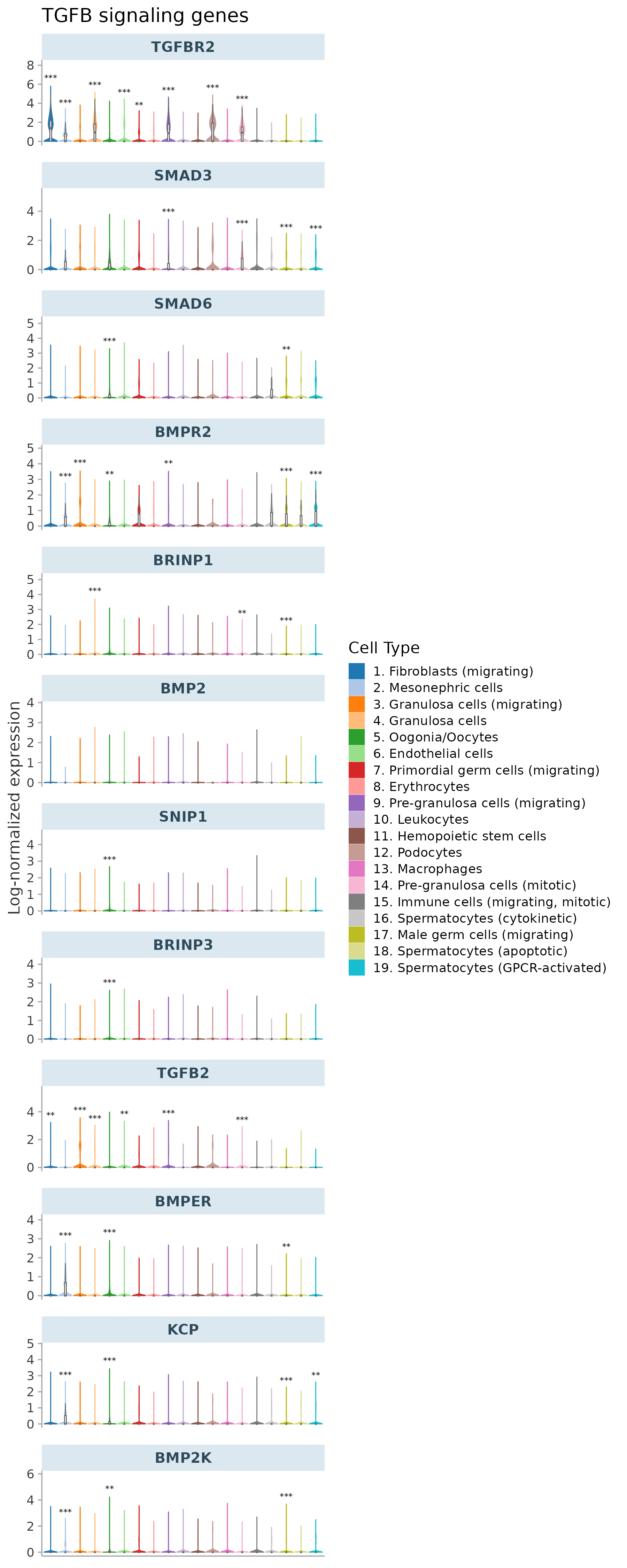

### Supplementary Figure S20

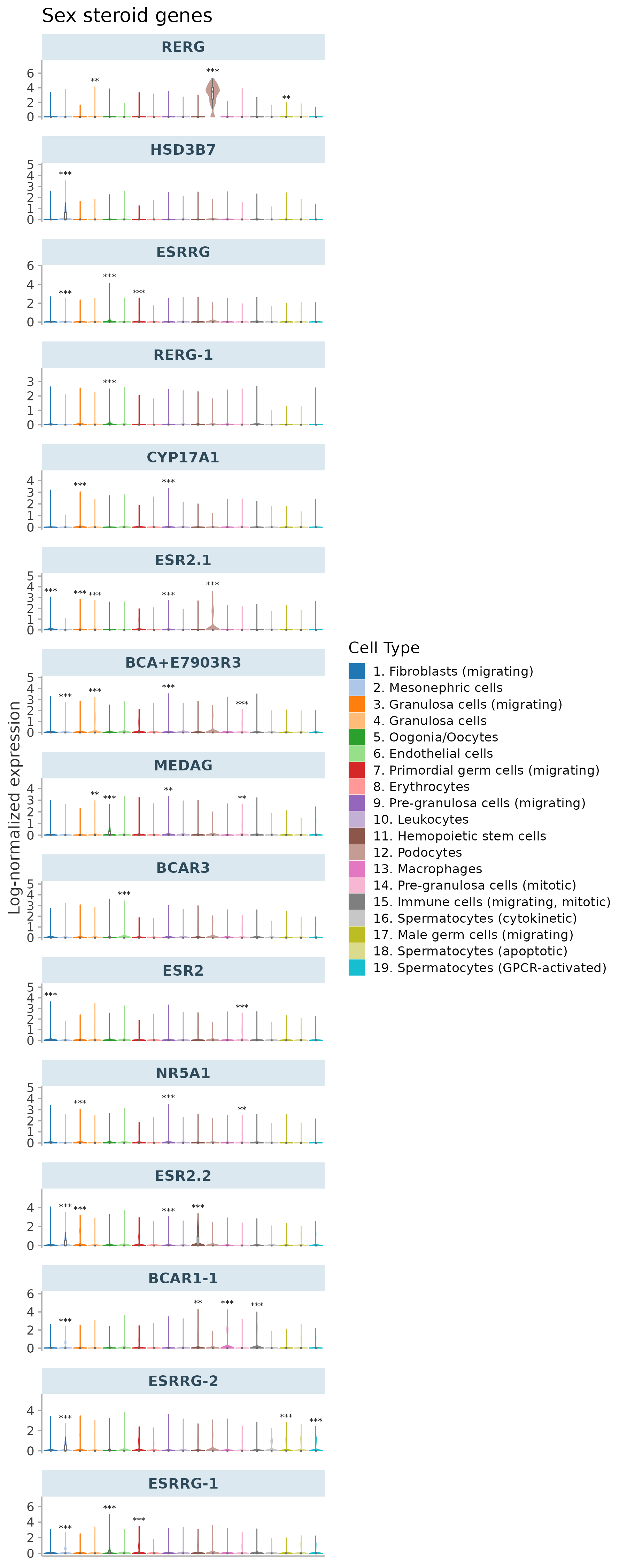

### Supplementary Figure S21

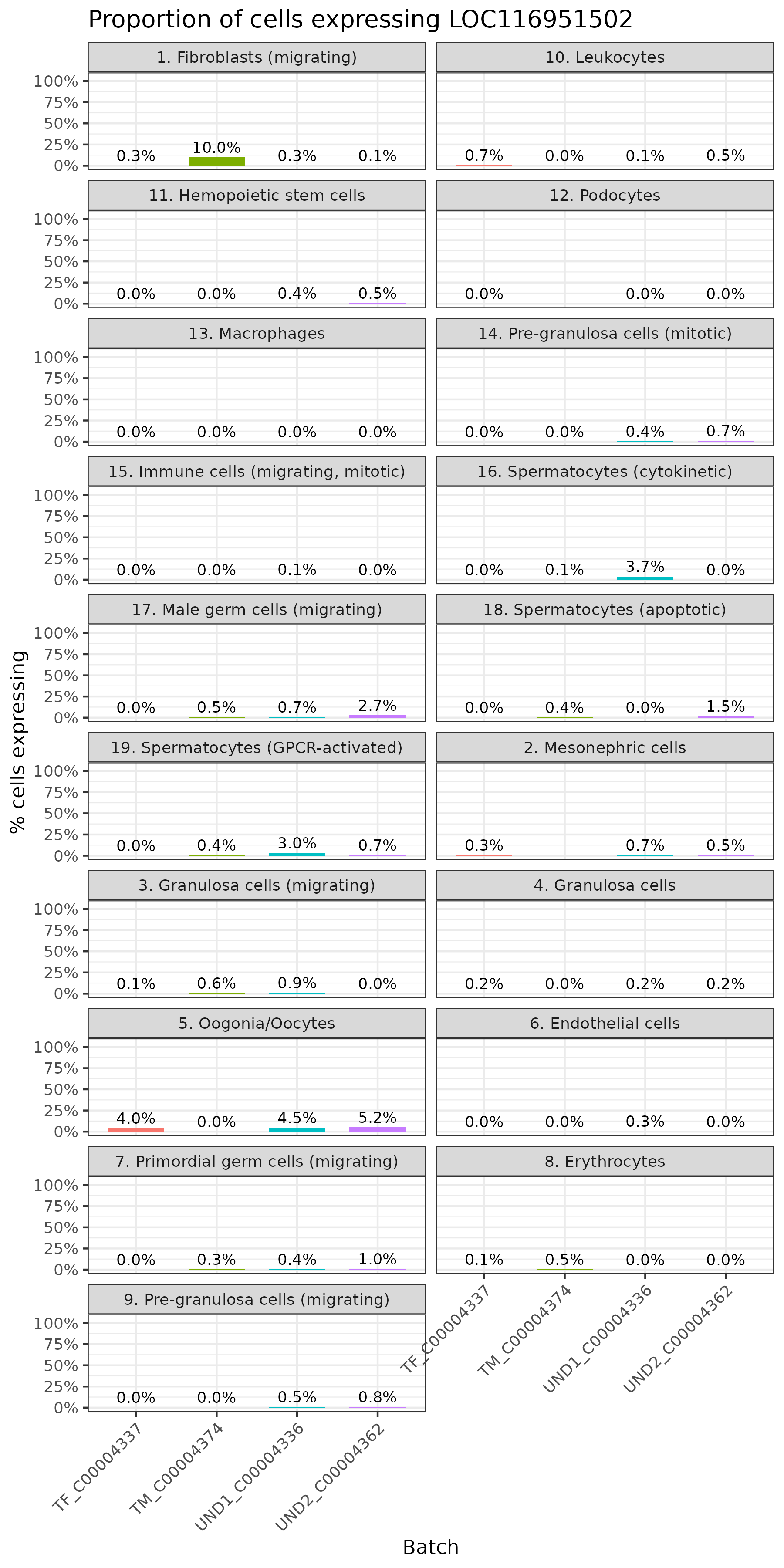

### Supplementary Figure S22

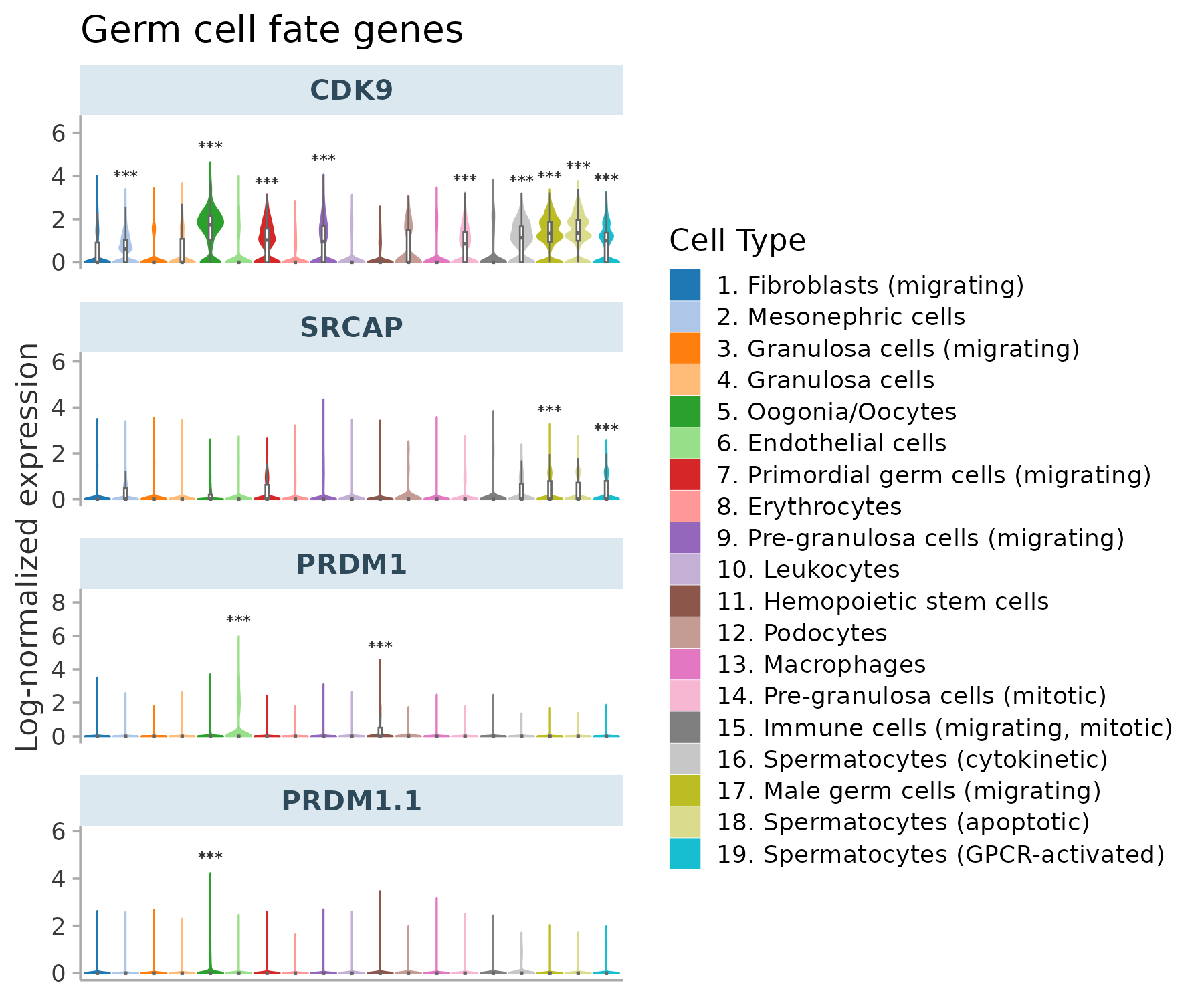
