## Supplementary Figure S10 for "Gonadal PIP-seq reveals genes involved in germ cell development and sexual differentiation in sea lamprey (*Petromyzon marinus*)"

### Slide 1
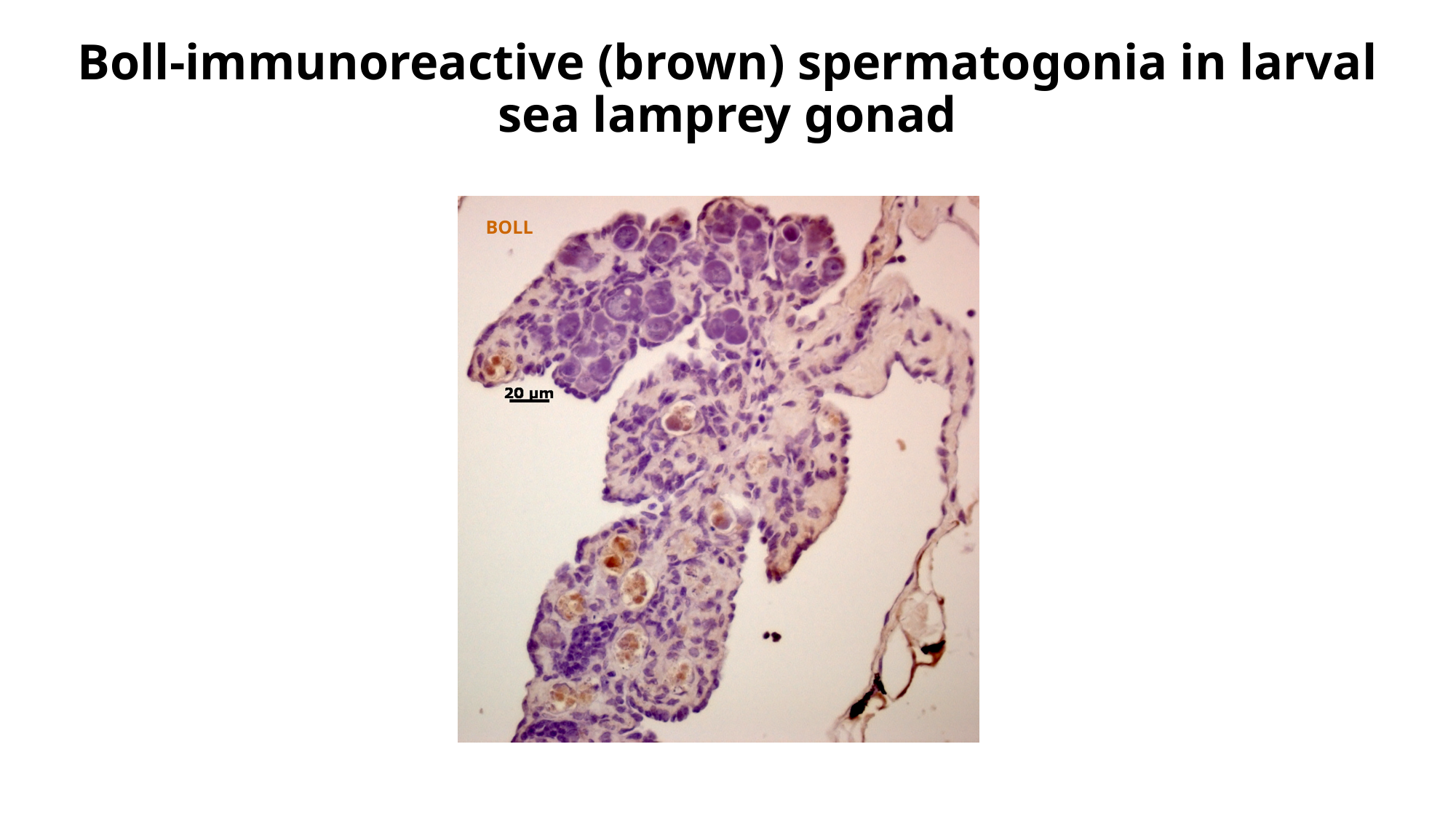

Boll-immunoreactive (brown) spermatogonia in larval sea lamprey gonad
BOLL

### Slide 2
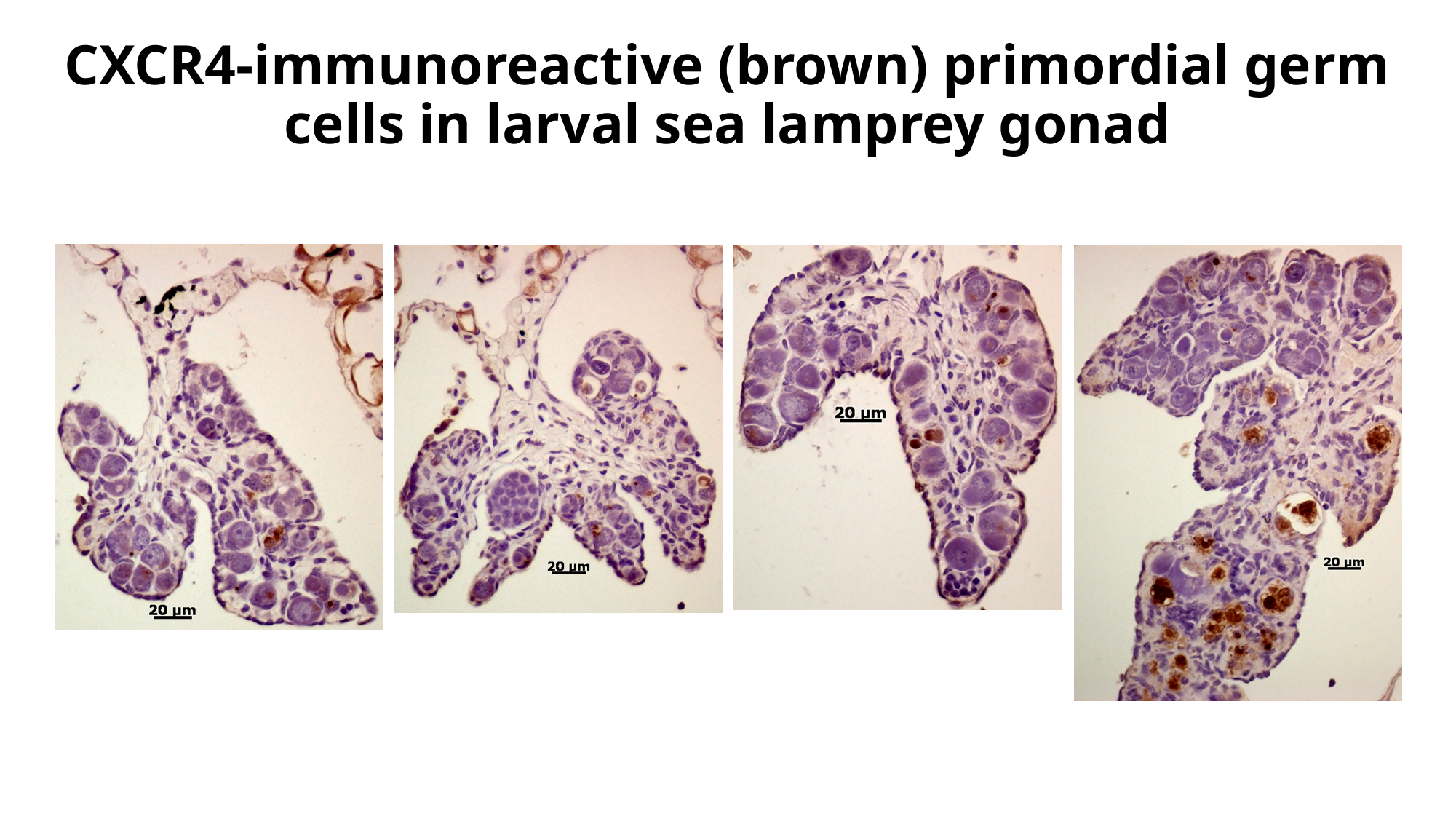

CXCR4-immunoreactive (brown) primordial germ cells in larval sea lamprey gonad

### Slide 3
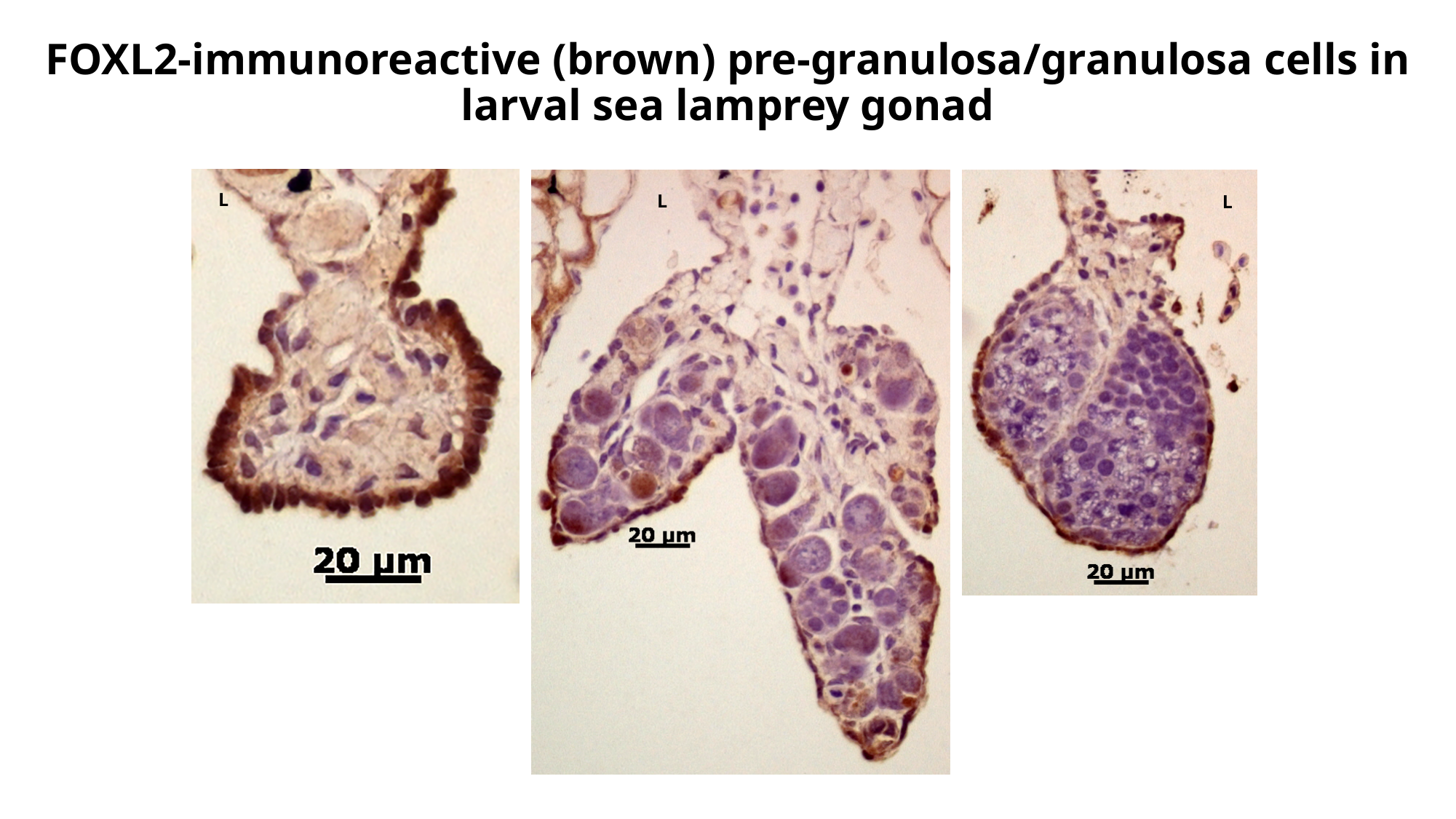

FOXL2-immunoreactive (brown) pre-granulosa/granulosa cells in larval sea lamprey gonad
L
L
L

### Slide 4
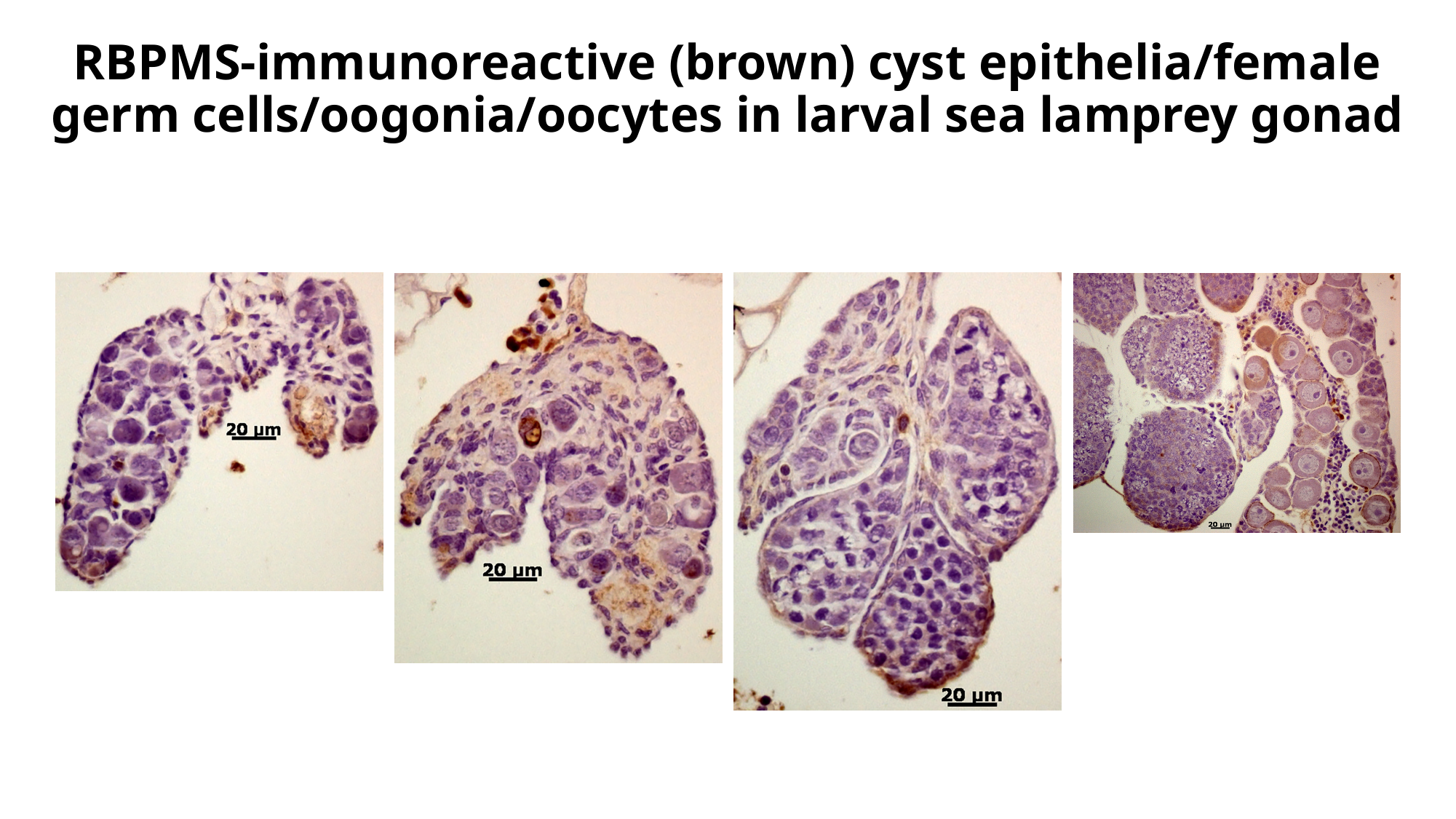

RBPMS-immunoreactive (brown) cyst epithelia/female germ cells/oogonia/oocytes in larval sea lamprey gonad

### Slide 5
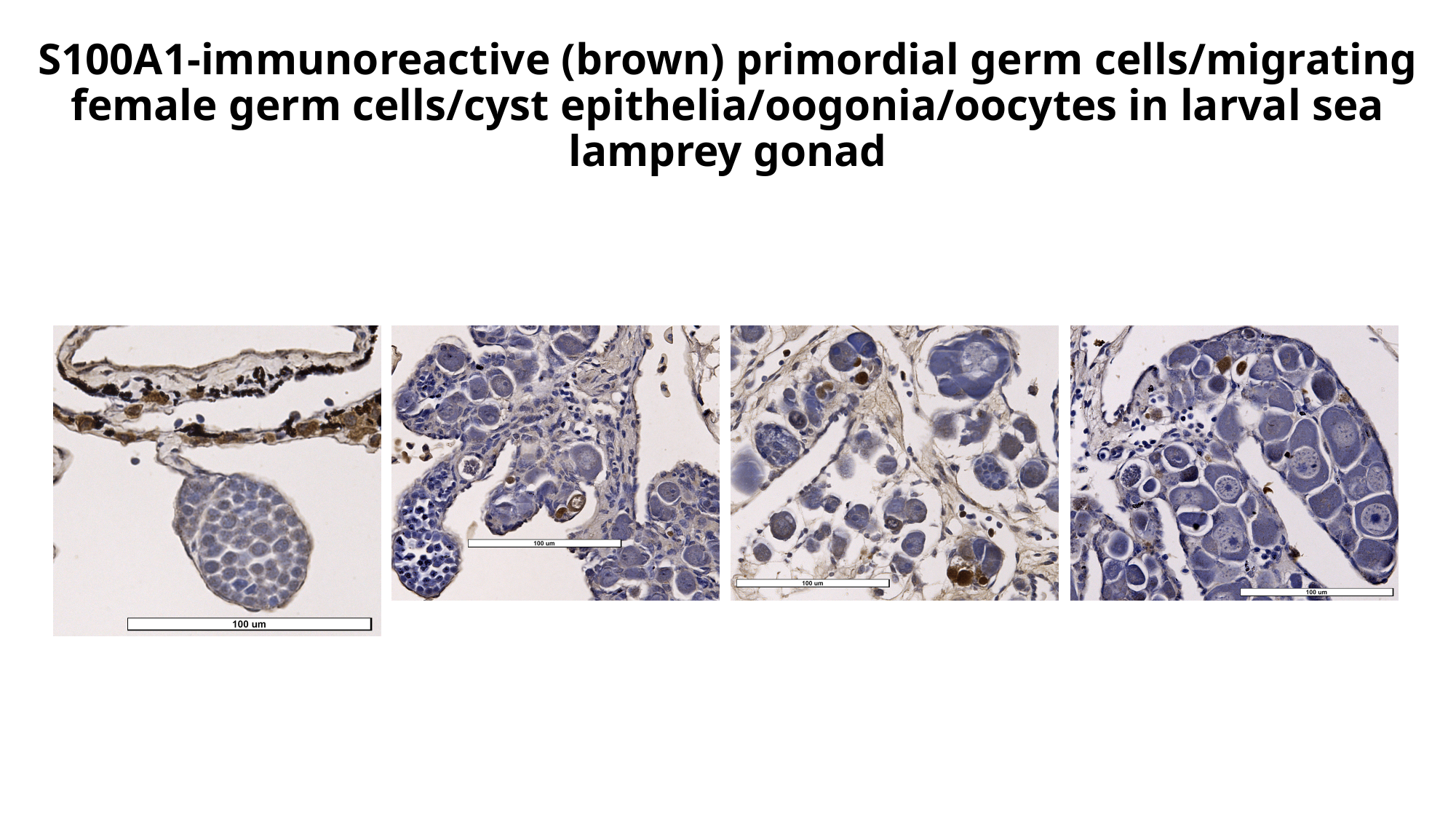

S100A1-immunoreactive (brown) primordial germ cells/migrating female germ cells/cyst epithelia/oogonia/oocytes in larval sea lamprey gonad

### Slide 6
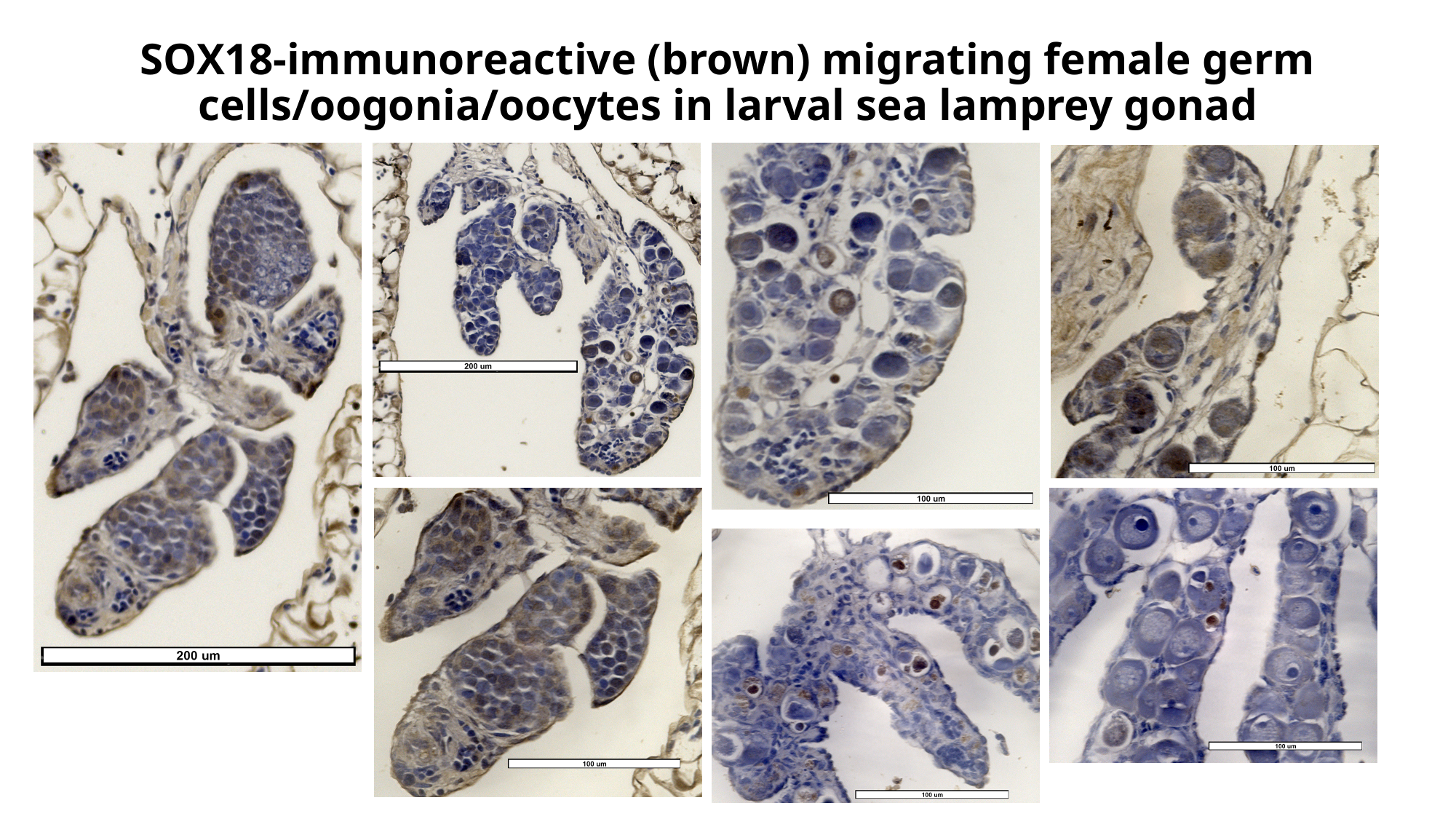

SOX18-immunoreactive (brown) migrating female germ cells/oogonia/oocytes in larval sea lamprey gonad
